# From Asymptomatic to Symptomatic: Multiomics profiling of the temporal response of grapevine viral-mixed infection

**DOI:** 10.1101/2023.07.17.549167

**Authors:** M.L. Fall, A. Poursalavati, A. Sidibé, D. Xu, P. Lemoyne, G.S. Martins, V.J. Javaran, P. Moffett, Carisse Odile

## Abstract

Mixed viral infections are common in grapevines. However, our understanding of the factors and signaling pathways that influence the expression of viral symptoms in mixed infections is still incomplete. In a previous study, we revealed that the presence of grapevine leafroll-associated virus species in mixed infections was randomly associated with the development of virus-like symptoms. To understand what drives the timing of these virus-like symptoms in mixed infections, we used dsRNA and total RNA sequencing and metabolomic analysis to profile the viromes, metabolites, and transcripts of grapevine leaves collected at two different times of the year (summer and autumn). We demonstrated that neither viral titre nor virome composition changes were associated with symptom expression in autumn. The total phenolic content and antioxidant capacity increased in most plants except for those with early onset symptoms. According to the results of differential gene expression analysis, cell wall biosynthesis pathways were significantly downregulated in all grapevine plants infected with grapevine leafroll-associated virus 3, grapevine asteroid mosaic-associated virus, and grapevine Pinot gris virus. In addition, polyketide pathways were significantly upregulated in all cultivars, while flavonoid precursor (e.g. abscisic acid) production was significantly reduced in plants that expressed strong virus-like symptoms. In the ‘Vidal’ cultivar, an uncharacterized double-stranded RNA-binding protein (DRB) appears to play a critical role in the plant’s antiviral defences, supporting the recent hypothesis that DRBs make an important contribution to dominant antiviral responses in plants. The seasonality of the expression virus-like symptoms appears to be a consequence of the dynamic interactions between antiviral factors and viral counter-defences that occur at different developmental stages of grapevine.

## INTRODUCTION

Grapevine is one of the most valuable crops worldwide, with over 7 million hectares planted in 2021, and a farm gate value of more than US$68 billion ^1, 2^. However, among cultivated crops, grapevine also hosts the largest number of viruses, including 87 species in 17 families and 34 genera ^3^. Viruses and viroids can decrease or impair rooting ability, graft take, root-stock-scion compatibility, overall vine vigour, yield, and fruit quality, resulting in major impacts to productivity. For decades, the identification and characterization of grapevine viruses has focused on taxa that cause visual symptoms and decrease yield. Advances in high-throughput sequencing over the past decade have revealed large numbers of cryptic, symptomless viruses and a very diverse grapevine virome (viruses and viroids) in term of composition, abundance, and the genetic variants within a given virus population ^4^. Some of these viruses, such as grapevine rupestris stem pitting-associated virus (GRSPaV), are believed to have evolved over many years of coexistence with grapevine ^5^. Indeed, grapevine is chronically infected by a number of viruses and virus-like particles that have little or no negative biological impact on their hosts. These viruses, which include GRSPaV, hop stunt viroid, grapevine yellow speckle viroid, and others form the so-called background virome. They often cause complex symptoms that are difficult to associate with a single causal agent ^6^. In the case of a large number of grapevines viruses (e.g. grapevine fleck virus, grapevine endophyte endornavirus), their association with virus-like symptoms has not been definitively established and needs further investigation because they are often present in co-infections with multiple other virus species ^7 4, 8^. In addition, the expression of symptoms can vary depending on the cultivar, and generally tends to be more pronounced during the cooler temperatures found in autumn ^9^. On the other hand, other virus species such as Arabis mosaic virus, grapevine fanleaf virus, grapevine Pinot gris virus, and grapevine leafroll-associated virus 1 and 3 cause distinctive symptoms in grapevine.

Grapevine leafroll-associated virus 3 (GLRaV3) and its genetic variants are the most wide-spread strains in grapevine and have been observed in grape production regions across the globe. This virus is known to cause well-described visual symptoms that include green veins and reddish interveinal areas in red cultivars and yellowing or chlorotic mottling in white cultivars ^7^. However, certain white cultivars such as ‘Vidal,’ ‘Thompson Seedless,’ and ‘Sauvignon Blanc’ express little or no symptoms when infected with GLRaV3 ^4, 10^. In addi-tion, in a recent study, our group found that the presence of various grapevine lea-froll-associated viruses is randomly associated with symptom expression ^4^. In addition, some genetic variants of GLRaV3 are known to produce no symptoms, while others cause well-described symptoms ^11^. Indeed, symptom expression and its intensity seem to be related to the strength of viral RNA-silencing suppressor activities, which differs among the phylogenetic groups of GLRaV3 ^12^.

Mixed virus and variant infections in the same plant are common in grapevine ^13^, causing complex, multifaceted host-virome interactions that influence symptom expression. The main objective of this study was to improve our understanding of the factors that drive the expression of virus-like symptoms in grapevine plants suffering from mixed infections. We aim to examine three hypotheses: (i) the composition and abundance of the grapevine virome changes over time; (ii) this temporal change in virome composition and abundance is associated with symptom expression in autumn; (iii) the temporal change in grapevines’ defence responses to viruses is associated with symptom expression.

Here we report that the virome composition and abundance were not significantly different when sampled in summer and in autumn and no association was found between symptom expression and the increased relative abundance of GLRaV3, grapevine asteroid mo-saic-associated virus (GAMaV), or grapevine Pinot gris virus (GPGV), all known to cause symptoms. This has major implications for grapevine virus diagnostics. Differential gene expression analysis revealed the downregulation of the genes involved in cell wall biosynthesis pathways in infected grapevine plants, and the upregulation of polyketide pathways in all cultivars.

## RESULTS

All the raw reads were submitted to the Sequence Read Archive (SRA): Bioproject PRJNA853579, biosamples from SAMN29398272 to SAMN29398307 (SRA from 29398272 to 29398307). All the data are available in the supplementary file.

### Grapevine virome profile

Overall, 83.33% of the grapevine plants (n =18) expressed virus-like symptoms in autumn, while only 11.11% of the plants did so in summer (Fig. 1 and Fig. S2). An average of 440,000 reads per sample were obtained with dsRNA sequencing, with viral reads making up between 2% and 20.5% of the reads. The viromes of 18 different grapevine plants were determined individually at two times in the growing season, summer and autumn. The positive control virus (PvEV1) was detected in all samples. Only viruses that were detected by both pipelines with a whole-genome coverage of at least 0.5 and a weight (titre) of greater than 0.001 were considered to be positive detections and hence suitable for downstream analysis. A total of 10 viruses and 2 viroids were identified, including grapevine asteroid mosaic-associated virus (GAMaV), grapevine fleck virus (GFkV), grapevine leafroll-associated virus 2 (GLRaV-2) and 3 (GLRaV-3), grapevine Pinot gris virus (GPGV), grapevine rupestris stem pit-ting-associated virus (GRSPaV), grapevine Syrah virus 1, grapevine virus B, grapevine virus E, grapevine virus H, grapevine yellow speckle viroid 1 (GYSVd 1), and hop stunt viroid (HSVd) (Fig. 2A). GRSPaV dominated the virome in most of the samples, followed by GYSVd 1, HSVd and GLRaV-3 in descending order. Mixed infections were predominant, with at least two viruses or viroids present in all sample (Fig. 2A). The number of species as a function of sampling effort reached a plateau after 12 samples, suggesting that viral species diversity was accurately captured and that most of the viral species were present in the majority of sampled plants (Fig. 2B). In 75% of the cases, the viral load (abundance of all detected viruses) did not change or decreased between the two sampling times (summer and autumn), despite the fact that 80% of symptomatic plants (n = 15) expressed symptoms only during the autumn. The relative abundance of GLRaV-3, which is known to cause notable symptoms, increased and decreased from summer to autumn in 11 % and 39% of infected plants respectively, while only one GLRaV-3-infected plant expressed symptoms during the summer (Fig. 2C). The relative abundance of GPGV, another virus known to cause symptoms, increased and decreased from summer to autumn in 33% and 28% of the GPGV-infected plants respectively. Three of the GPGV-infected plants expressed symptoms during the summer and the relative abundance of GPGV in these plants decreased between the two sampling times. One grapevine plant (CO1216) infected by both GLRaV-3 and GPGV showed no symptoms in either summer or autumn, even though the relative abundance of these viruses increased between the two sampling times (Fig. 2C). However, according to the non-parametric and dissimilarity tests, no significant (P < 0.001) difference was found in viral abundance (all viruses considered) at the two sampling times in all samples (Table 1, Fig. 3A-B). The three different community diversity indices, and particularly the Morista-Horn index, had values between 0.95 and 1, except for one sample, DSJPSB (M = 0.77) (Table 2, Fig. 3C-H). The viral communities obtained at the two different sampling times were highly similar in species richness, composition and abundance (Table 2, Fig. 3C-H).

**FIG 1.**
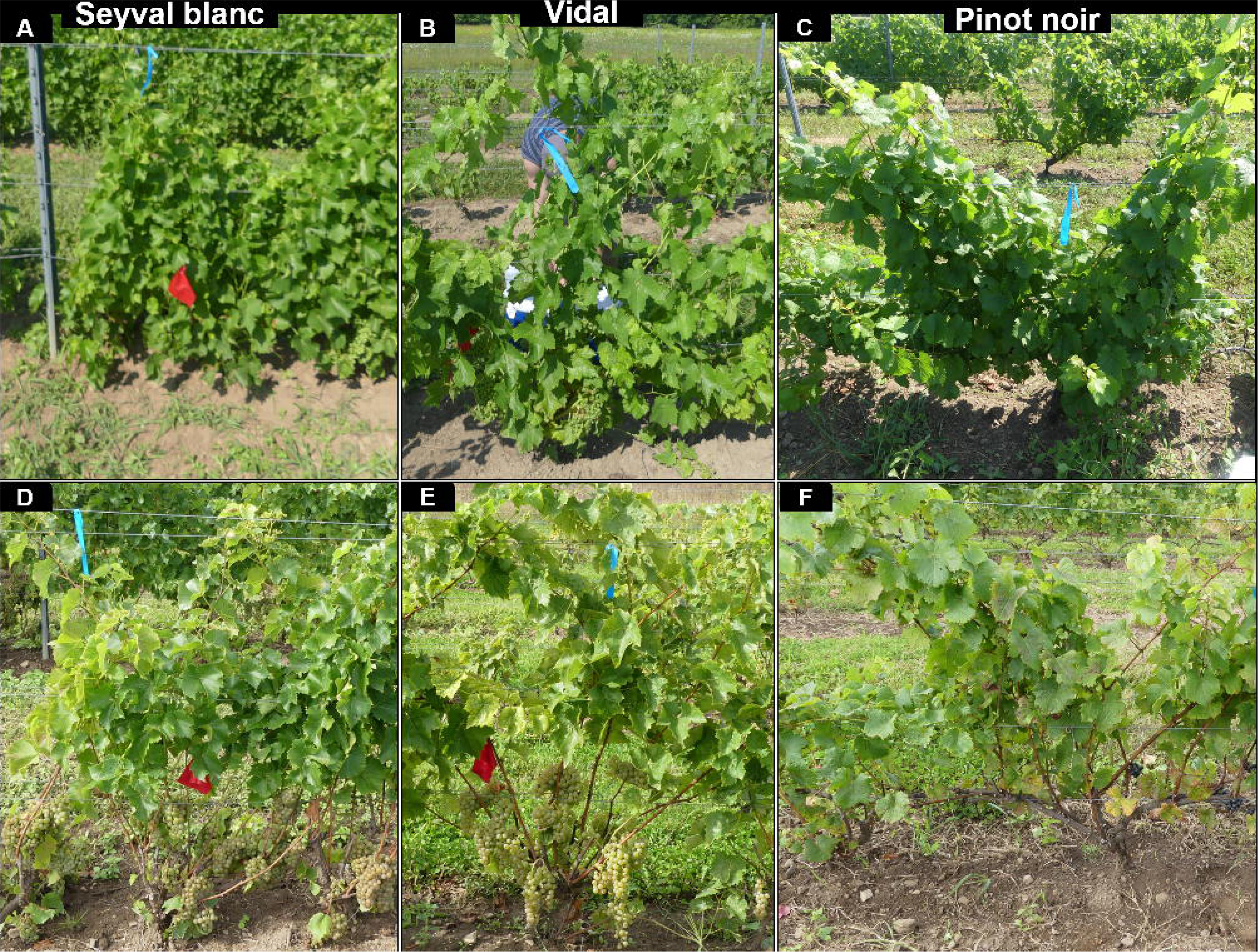
Symptomatology of grapevine in leaves collected at Farm 1 at two sampling times. **A** represents plant BacSB (‘Seyval Blanc’), **B** represents plant BacV3 (‘Vidal’), and **C** represents plant BacPN1 (‘Pinot Noir’), samples of which were collected during the summer**. D** repre-sents plant DSJSB5 (‘Seyval Blanc’), **E** represents plant DSJV5 (‘Vidal’), and F represents plant BacPN4 (‘Pinot Noir’), samples of which were collected during the autumn. The virome, transcriptome and metabolome profiles of these grapevine plants were determined.

**FIG 2.**
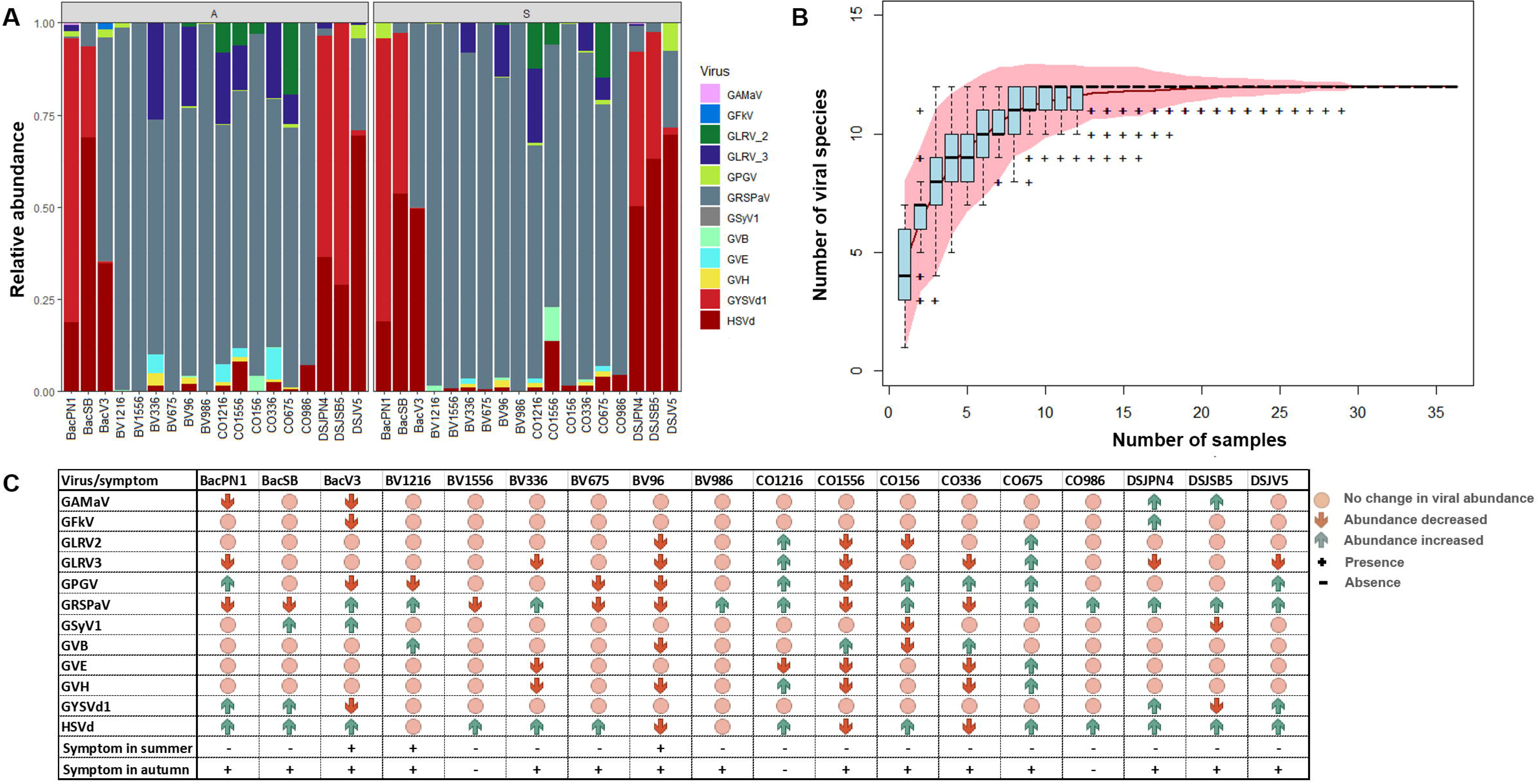
Temporal dynamics of viral species in samples from 18 selected grapevine plants col-lected at two times during the growing season. **A**, normalized relative abundance of viral spe-cies during the autumn (A, 1st panel) and the summer (S, 2nd panel). **B**, accumulation curve of viral species as a function of the number of samples. The red line represents the number of viral species as a function of the number of samples collected. The pink area displays the standard deviation, and the blue box plots show the species richness based on the linear interpolation of random permutations. **C**, heat table displaying the dynamics of viral species abundance by sampling time as a function of virus-like symptom status (asymptomatic, symptomatic).

**FIG 3.**
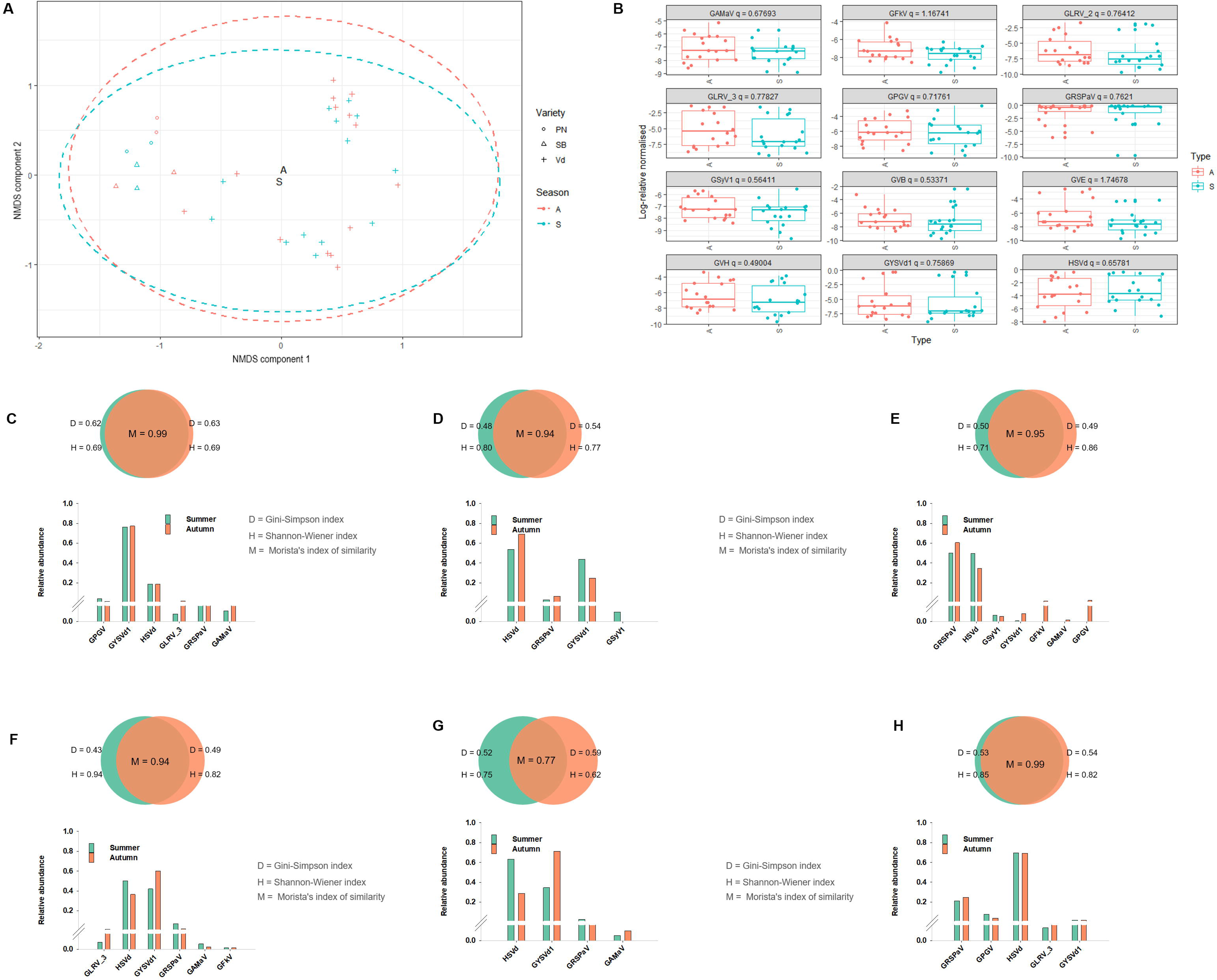
Comparison of the viral abundance in samples from 18 selected grapevine plants col-lected at two times (summer vs. autumn) during the growing season. **A**, non-metric multidi-mensional scaling (NMDS) plot displaying the dissimilarities in viral abundance between the two sampling times. A Bray-Curtis dissimilarity matrix was generated and the covariance matrix was calculated (Buttigieg and Ramette 2014; Ramette 2007). **B**, graphical representa-tion of the non-parametric test (Kruskal-Wallis) using the kruskal.test function iteratively, with the value of alpha set at 0.01. The p-value, the number of false positives expected by chance (E-value), and the probability of finding by chance at least one false positive were calculated (family wise error rate [FWER]). A significant difference in viral abundance at the two sampling times was defined as one in which the q-value, which is the adjusted p-value after the Benjamini and Hochberg correction, is less than 0.01 (Benjamini and Hochberg 1995; Storey 2003). The relative abundance of viral species in selected grapevine plants at Farm 1 and Farm 2 is also shown; **C** represents plant BacSB (‘Seyval Blanc’) at Farm 1, **D** represents plant BacV3 (‘Vidal’) at Farm 1, **E** represents plant BacPN1 (‘Pinot Noir’) at Farm 1, **F** represents plant DSJSB5 (‘Seyval Blanc’) at Farm 2, **G** represents plant DSJV5 (‘Vidal’) at Farm 2, and **H** represents plant BacPN4 (‘Pinot Noir’) at Farm 2. The virome, transcriptome and metabolome profiles of these grapevine plants were determined. The Gini-Simpson index (D) and the Shannon-Wiener diversity index (H) were calculated for each sampling time and Morista’s index of similarity (M) was calculated to measure the similarity of the viral communities (species richness and abundance) obtained at the two sampling times.

**Table 1.**
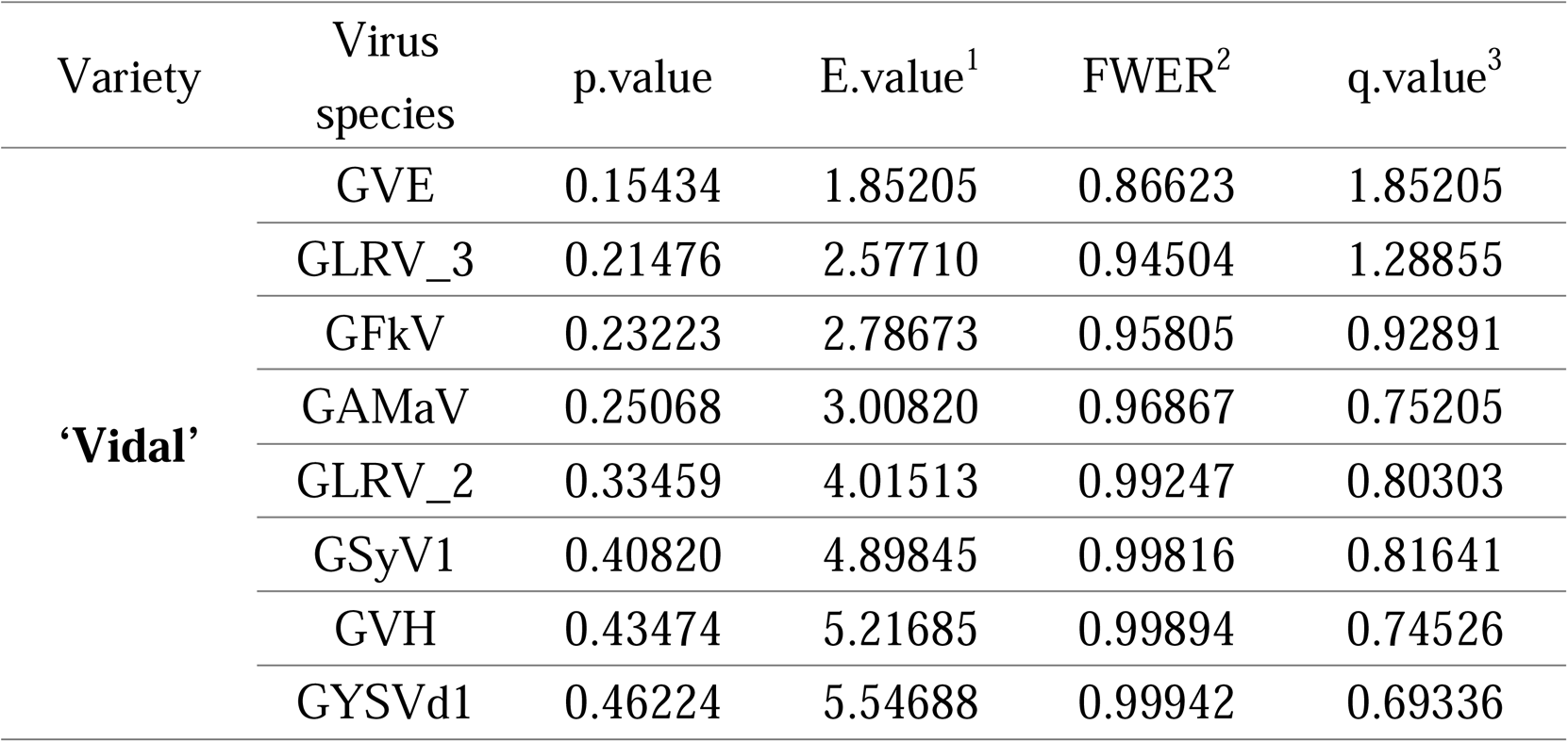
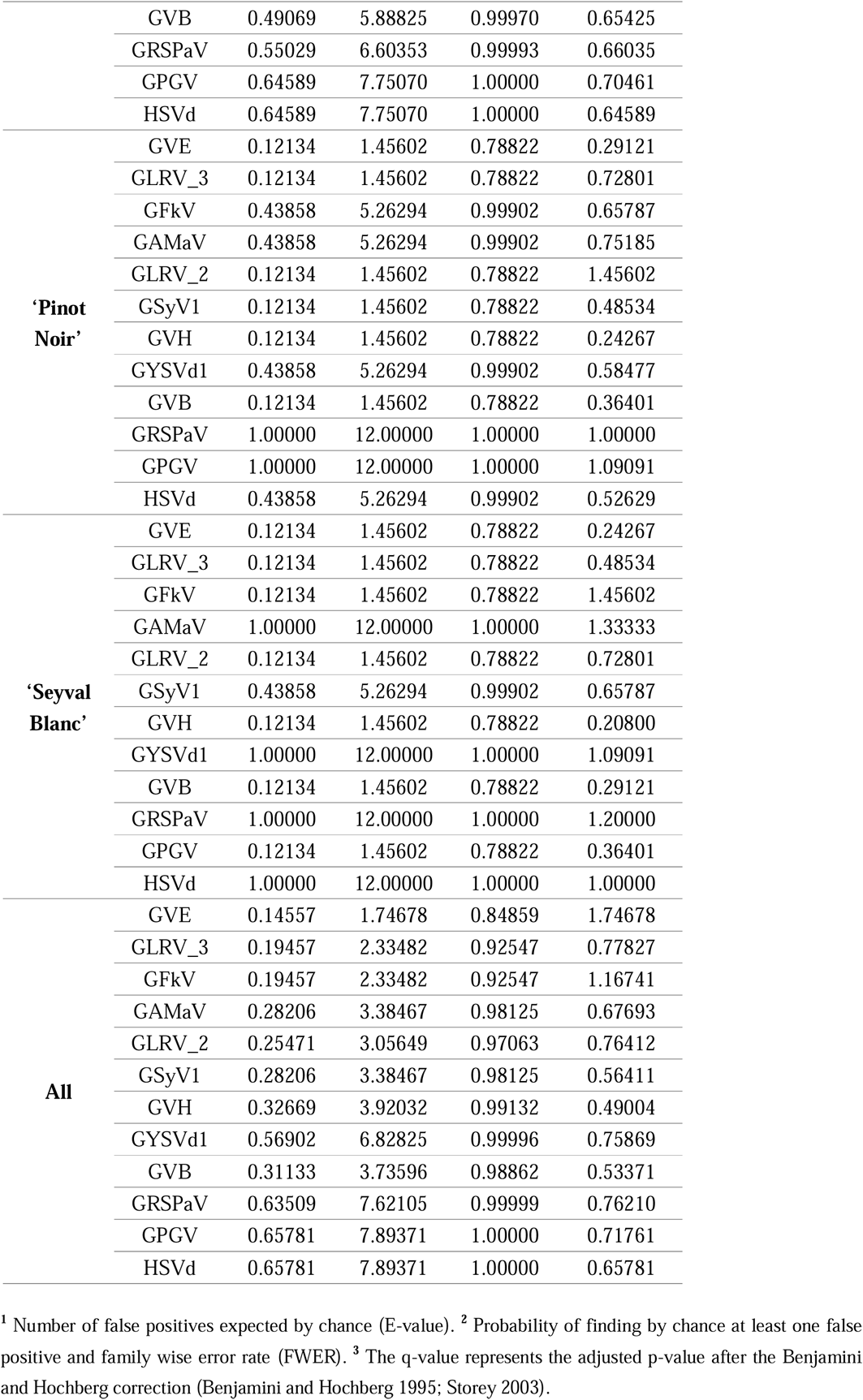
Non-parametric comparative analysis of viral abundance at the two sampling times (summer vs. autumn). An E-value much smaller than 1 indicates a significant difference in viral abundance between the two sampling points. The q-value is the adjusted p-value after the Benjamini and Hochberg correction.

**Table 2.**
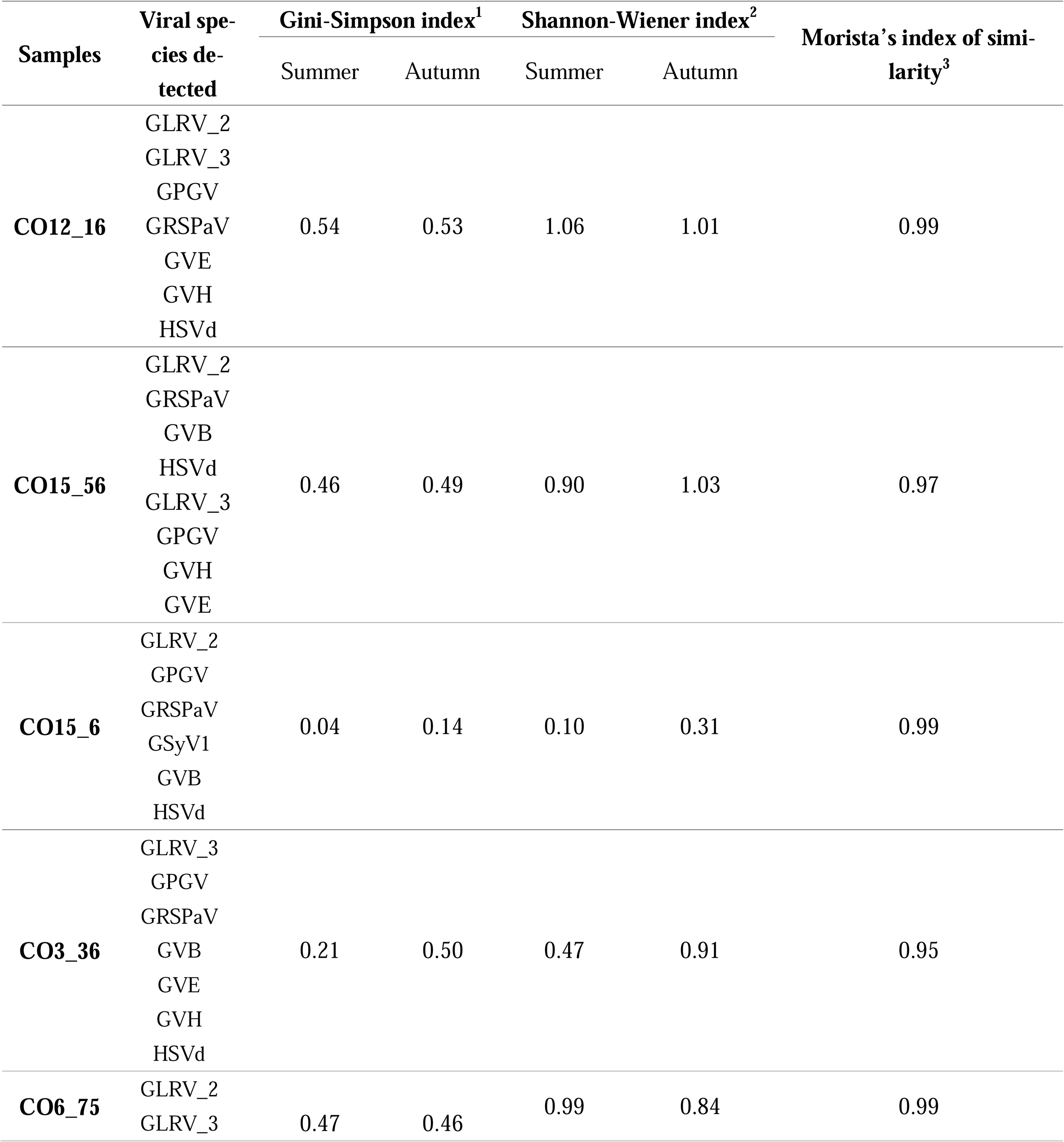
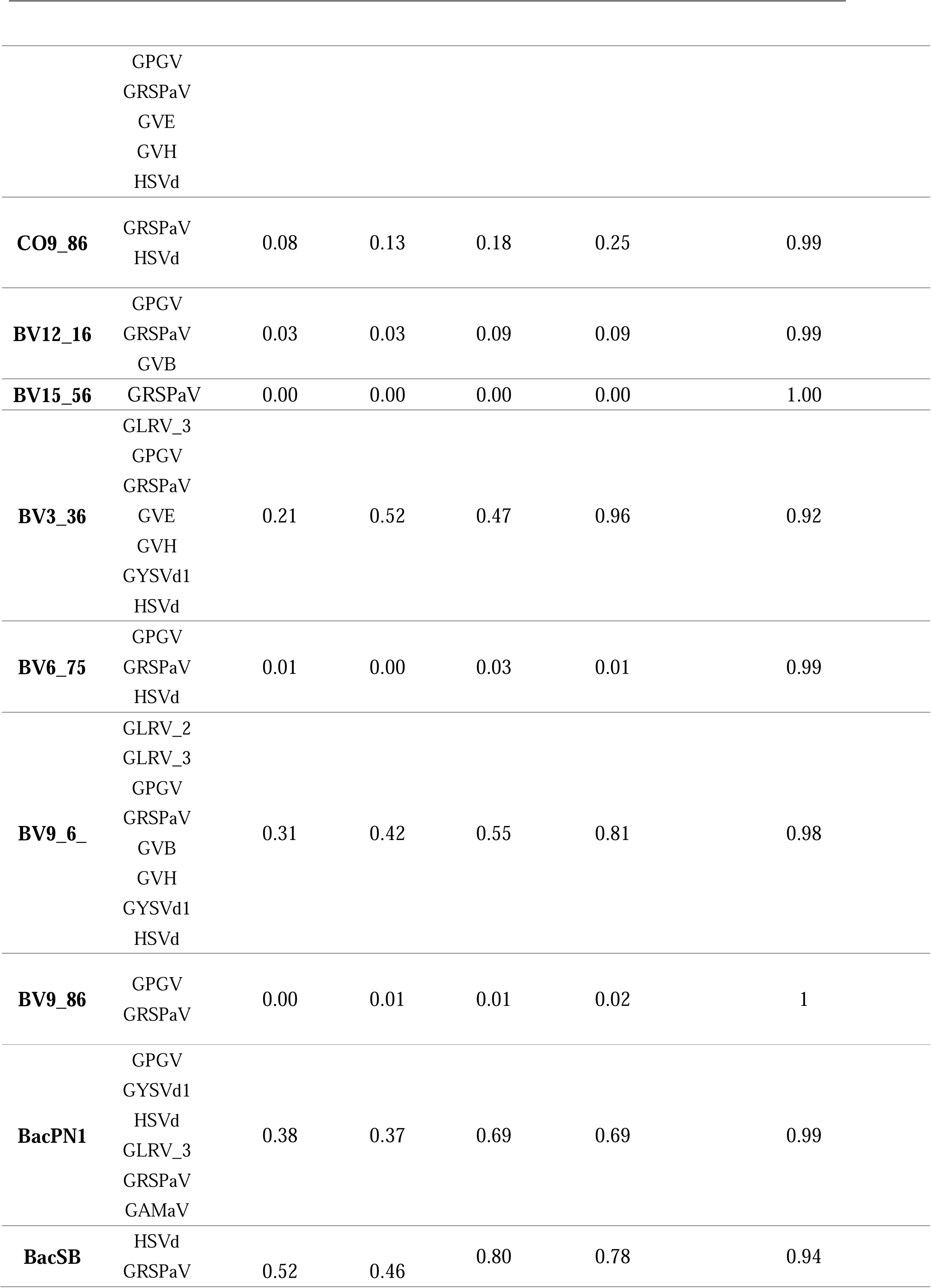
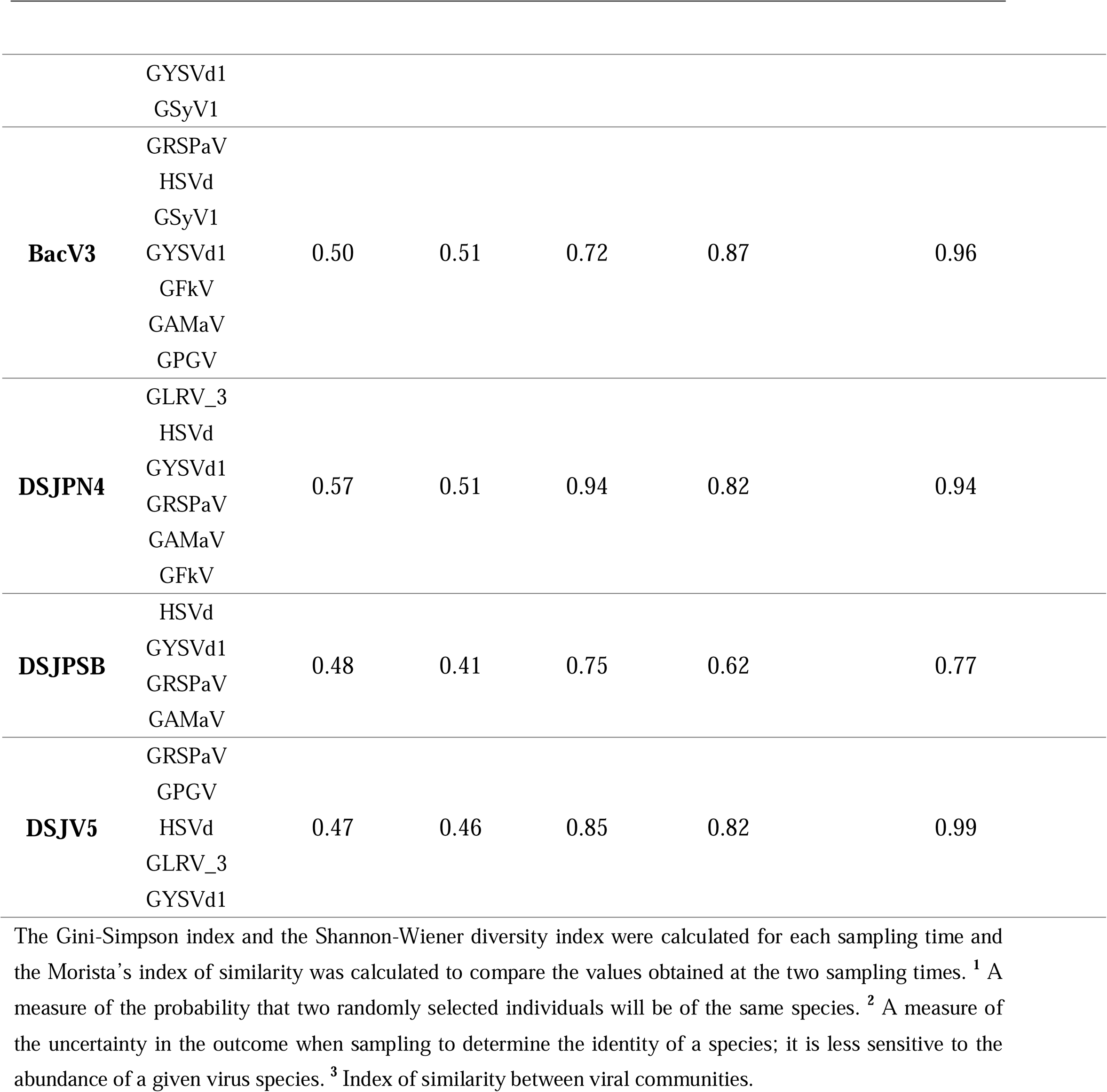
Temporal dynamics of the similarity in viral diversity and viral community at the two sampling times. Three different community diversity indices were calculated and are shown in this table.

### Differential gene expression (DGE) profile for host and viruses

The total RNA sequencing of the selected plants expressing virus-like symptoms yielded an average of 89 million reads, with the proportion of mapped viral reads ranging between 0.07% and 0.14%. The selected plants (12 samples) used for DGE analysis expressed different levels of symptoms during the autumn, despite the fact that the viral communities were highly similar in between the two sampling times in term of species richness, composition and abundance (Fig 1, Fig. S2, Table 2, Fig. 3C-H). Virus-like symptoms were more severe in the ‘Pinot Noir’ cultivar than in the ‘Vidal’ and ‘Seyval Blanc’ cultivars (Fig. 1, Fig. S2).

#### Grapevine DGE profile

##### DGE profile for metabolite processes

When the cultivar effect is not considered, a total of 27,642 genes were expressed and only 833 and 1,011 genes were expressed exclusively in the autumn and summer, respectively. A total of 3,539 and 3,995 grapevine-related genes underwent significant upregulation and downregulation, respectively (Fig. S3). Regardless of the cultivar, the main upregulated pathways involved aromatic polyketide metabolic pathways (e.g. stilbenoid, and gingerol biosynthesis), cellular carbohydrate catabolic processes, circadian rhythm regulation (e.g. cold response processes), galactose metabolism, and others. The main downregulated pathways included anion antiporter activity (e.g. inorganic anion transmembrane transporter and sulfate compound transporter activity), responses to light stimulus and cell wall biosynthesis (photosynthesis, response to radiation, etc.), fatty acid biosynthesis, nitrogen metabolism, and pectin catabolic processes (Fig. S3).

##### In ‘Seyval Blanc’ (SB)

In the samples from Farm 1, the main upregulated pathways involved iron homeostasis (43%), aromatic polyketide metabolism (12%), cell wall biosynthesis (30%), and circadian rhythm regulation (10%), while downregulated pathways included isoprenoid metabolism (46%), exopeptidase activity (30%), and steroid and fatty acid biosynthesis (23%). In contrast, in the samples from Farm 2, the main upregulated pathways were related to circadian rhythm regulation (50%), aromatic polyketide metabolism (29%) and phenylpropanoid biosynthesis (16%), while downregulated pathways involved cell wall biosynthesis (40%), temperature responses (34%), and endopeptidase processes (9%) (Fig. S4).

##### In ‘Pinot Noir’ (Pn)

In the Farm 1 samples, the main upregulated pathways involved phenylpropanoid biosynthesis (70%) and polyketide metabolism (23%), while the down-regulated ones included cell wall biosynthesis (21%), carboxypeptidase activity (14%), and isoprenoid biosynthesis (10%). In the Farm 2 samples, the main upregulated pathways were related to iron homeostasis (45%), phenylpropanoid biosynthesis (19%), and polyketide metabolism (23%), while the downregulated pathways involved cell wall biosynthesis (69%), isoprenoid biosynthesis (12%), and phloem or xylem histogenesis (6%) (Fig. S4).

##### In ‘Vidal Blanc’ (V)

In the Farm 1 samples, the main upregulated pathways were related to DNA-binding transcription activity (42%) and aromatic polyketide metabolism (21%), and the downregulated pathways involved cell wall biosynthesis (49%), polysaccharide metabolic processes (10%), and sterol biosynthesis (9%). In the Farm 2 samples, the main upregulated pathways included polyketide metabolism (35%) and circadian rhythm regulation (5%) and the downregulated pathways, cell wall biosynthesis (29%), isoprenoid biosynthesis (12%), and sterol biosynthesis (20%) (Fig. S4).

##### DGE profile for RNAi-related genes

A total of 65 different genes involved in RNAi pathways exhibited significant differential regulation between the two sampling times. Regardless of cultivar, 31 genes were expressed differentially (six upregulated and 25 downregulated), with 37% of the significantly downregulated genes related to grapevine’s defence responses to viral infection (Fig. 4). Genes exhibiting significant regulation against viral infection include those coding for RNA-dependent RNA polymerase (RDR) 1, 2, 5, and 6 (VIT_01s0011g05880, VIT_17s0000g07980, VIT_11s0016g03220, and VIT_04s0008g05430), endoribonuclease Dicer homolog (DCL) 1, 2, and 4 (VIT_15s0048g02380, VIT_04s0023g00920, and VIT_19s0090g00940), double-stranded RNA binding protein (VIT_14s0006g01440), THO complex protein (VIT_11s0016g01910), Argonaute 1, 2, 4, 5, 9, and 10 (VIT_05s0020g04190, VIT_10s0042g01200, VIT_06s0009g01200, VIT_06s0061g01040, VIT_06s0009g01200, and VIT_05s0020g04190), RNA helicase-like protein (SDE3) (VIT_05s0020g03760), and others (Fig. 4). Among the cases of DGE involving RNAi-related genes, 22 (Dicer-like proteins, Argonaute proteins, RNA-dependent RNA polymerase) were validated by quantitative PCR (Figs. 5A and B).

**FIG 4.**
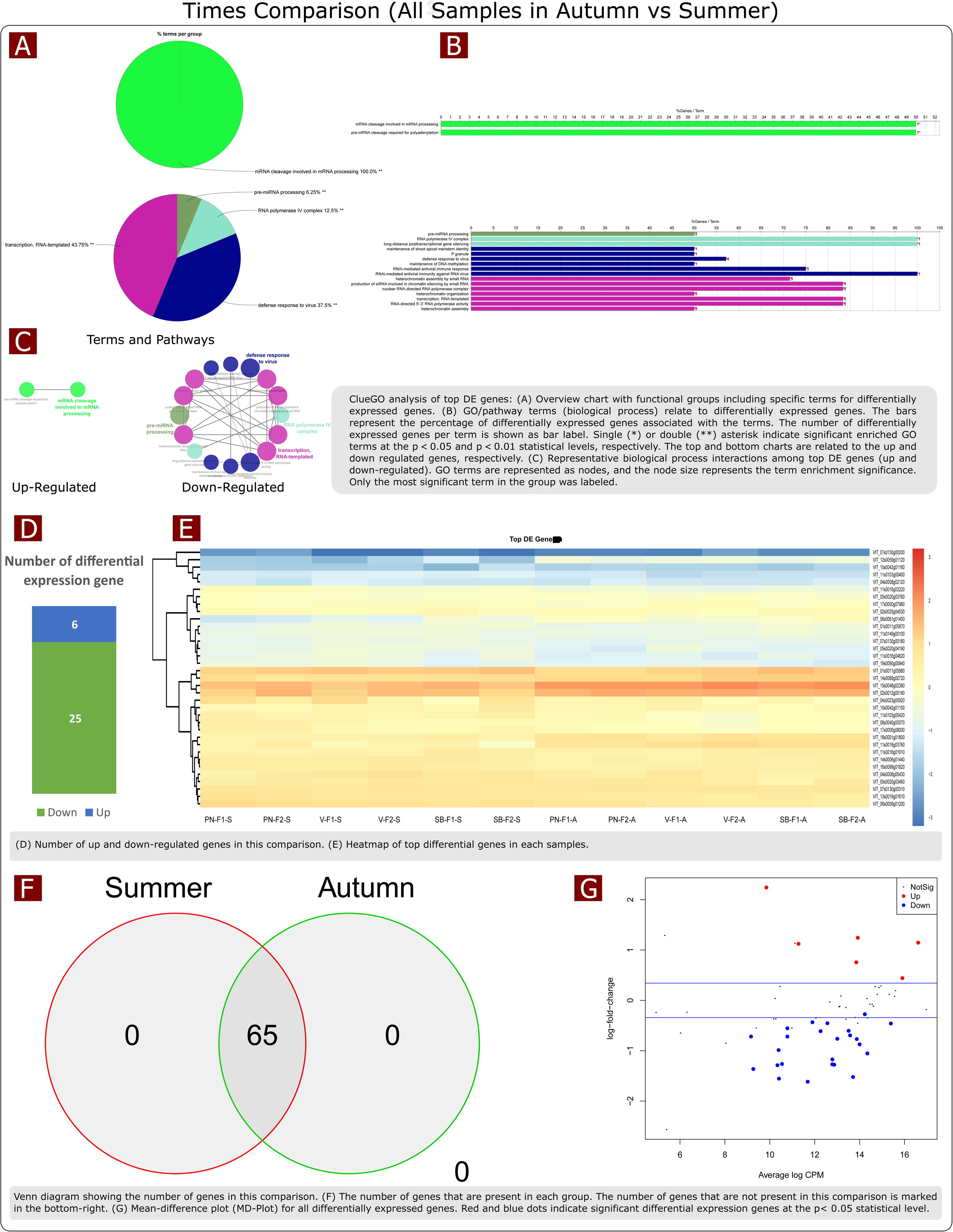
Graphical representations of functional gene groups related to RNA-silencing path-ways in grapevines differentially expressed at the two sampling times (summer vs. autumn) without consideration of cultivars. **A**, overview chart with functional groups including spe-cific terms for differentially expressed genes. **B**, gene ontology (GO)/pathway terms (bio-logical processes) relating to differentially expressed genes. The bars represent the percentage of differentially expressed genes associated with the terms. The number of differentially ex-pressed genes per term is shown on the bar label. Single (*) or double (**) asterisks indicate significantly enriched GO terms at the p < 0.05 and p < 0.01 statistical levels, respectively. The top and bottom charts show upregulated and downregulated genes, respectively. **C**, graphical representation of interactions among the most significant differentially expressed genes (upregulated and downregulated) in biological processes. GO terms are represented as nodes, and the node size represents the enrichment significance of the term. Only the most significant term in the group was represented. **D**, number of upregulated and downregulated genes in the comparison. **E**, heatmap of most significant differentially expressed genes in each sample. **F**, number of genes present at each sampling time. The number of genes that are not present in this comparison is shown on the bottom right-hand side. **G**, mean-difference plot (MD-Plot) for all differentially expressed genes. Red and blue dots indicate genes with significant differential expression at the p < 0.05 statistical level.

**FIG 5.**
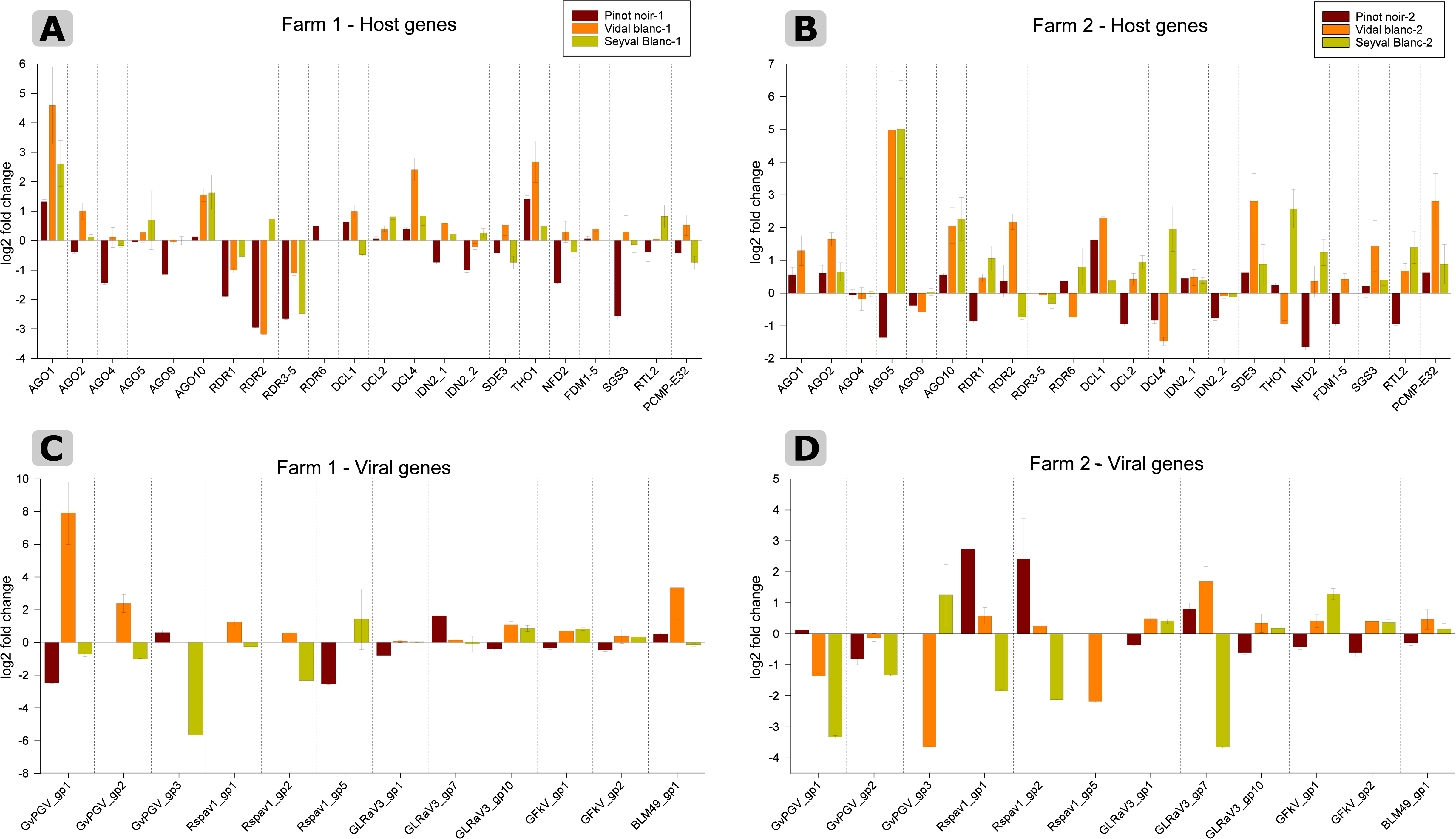
Quantitative RT-PCR validation of differential transcript expression analyses of 22 selected grapevine genes (e.g. Dicer-like proteins, Argonaute proteins, RNA-dependent RNA polymerase), **A** at Farm 1, **B** at Farm 2, with 12 viral genes (e.g. viral replicases, viral RNA silencing suppressors), **C** at Farm 1, **D** at Farm 2, which were detected by RNA sequencing. The histograms represent the transcript log2 fold change normalized to the control genes; the error bars refer to ± SE of three replicates.

##### In ‘Seyval Blanc’ (SB)

14 genes involved in RNAi pathways were significantly regulated (6 upregulated and 8 downregulated), with 42% of downregulated genes related to defence responses by grapevine to viral infection. The significantly (p < 0.01) downregulated genes included DCL 2, AGO2, AGO9, AGO10, RDR6, RDR3, and the grapevine homolog of the Arabidopsis factor of DNA methylation (FDM 1) (VIT_11s0103g00420), while significantly (p < 0.01) upregulated genes included SDE3, DCL1, a RNase III (VIT_12s0059g01120), and the grapevine homolog of the Arabidopsis SGS3 protein (VIT_07s0130g00190) (Fig. 6-1). In the Farm 1 samples, six additional genes, including DLC 3 (VIT_04s0008g02170) and DLC2, were significantly (p < 0.01) upregulated. A total of nine additional genes displayed significant downregulation, including the grapevine homolog of the Arabidopsis SERRATE (SE) gene (VIT_06s0009g01570) and RDR2 (VIT_17s0000g07980); in addition, 100% of RNA-directed RNA polymerase complex processing was downregulated. In the Farm 2 samples, four additional genes were upregulated, including THO complex subunit 3 (THO3) (VIT_19s0015g01220), SICKLE plant protein (VIT_18s0001g07660), and RTL2 (VIT_07s0129g00400), while eight genes were downregulated, including AGO10 (VIT_11s0016g04620), an uncharacterized RTL (VIT_17s0000g08000), the grapevine homolog of the Arabidopsis RNase III-like protein 2 (RTL2) (VIT_19s0090g00940), grapevine factor of DNA methylation (VIT_11s0103g00400), an uncharacterized AGO protein dissimilar to all other proteins (50% identity threshold) (VIT_06s0061g01040), an uncharacterized protein dissimilar to all other proteins (50% identity threshold) (VIT_10s0042g01160), an uncharacterized Argonaute-like protein 50% identical to AGO4 (VIT_08s0040g00070), and AGO2. In addition, 37% of the downregulated functional groups involved virus defence responses and 62%, RNA gene silencing (Fig. S5-1, supplementary file).

**FIG 6.**
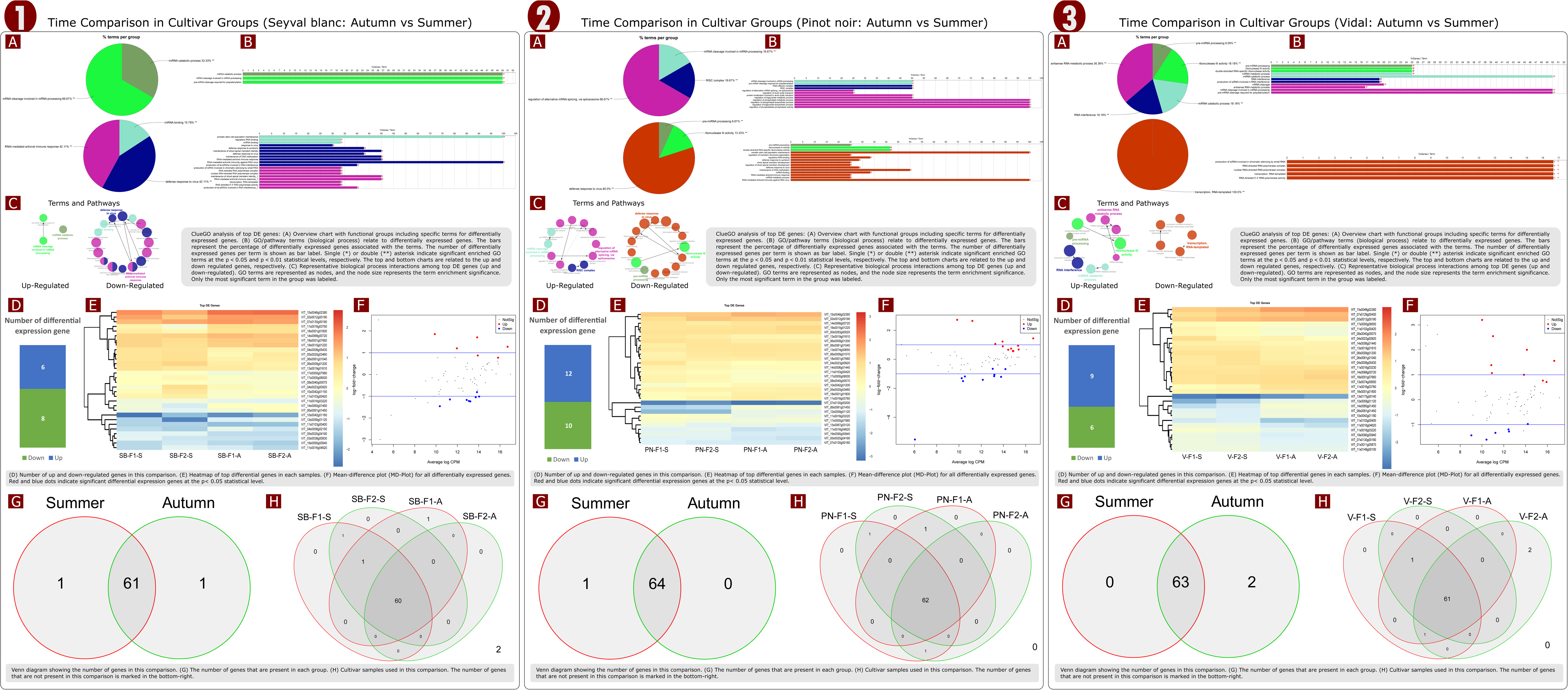
Graphical representations of functional gene groups related to the RNA-silencing pathways in grapevines and differentially expressed at the two sampling times (summer vs. autumn) grouped by cultivar, without consideration of the farm location. **Panel 1** represents ‘Seyval Blanc,’ **Panel 2** represents ‘Pinot Noir,’ and **Panel 3** represents ‘Vidal.’ **A**, overview chart with functional groups, including specific terms for differentially expressed genes. **B**, gene ontology (GO)/pathway terms (biological processes) relating to differentially expressed genes. The bars represent the percentage of differentially expressed genes associated with the terms. The number of differentially expressed genes per term is shown on the bar label. Sin-gle (*) or double (**) asterisks indicate significantly enriched GO terms at the p < 0.05 and p < 0.01 statistical levels, respectively. The top and bottom charts show upregulated and downregulated genes, respectively. **C**, graphical representation of interactions among the most significant differentially expressed genes (upregulated and downregulated) in biological processes. GO terms are represented as nodes, and the node size represents the enrichment significance of the term. Only the most significant term in the group was represented. **D**, number of upregulated and downregulated genes in the comparison. **E**, heatmap of most sig-nificant differentially expressed genes in each sample. **F**, mean-difference plot (MD-Plot) for all differentially expressed genes. Red and blue dots indicate genes with significant differen-tial expression at the p < 0.05 statistical level. **G**, Venn diagram of number of genes present at each sampling time. **H**, Venn diagram of number of genes present at each sampling time (S, summer, A, autumn) in samples at Farm 1 (F1) and Farm 2 (F2). The number of genes that are not present in this comparison is shown on the bottom right-hand side.

##### In ‘Pinot Noir’ (Pn)

22 genes involved in RNAi pathways were significantly regulated (12 upregulated and 10 downregulated), with 80% of downregulated genes associated with defence responses by grapevine to viral infection. Significantly (P < 0.01) downregulated genes included DCL 2, RDR3, AG10, the grapevine homolog of the Arabidopsis RNase III-like protein 2 (RTL2) (VIT_19s0090g00940), and the grapevine homolog of the Arabidopsis factor of DNA methylation (FDM 1) (VIT_11s0103g00420), while significantly (P < 0.01) upregulated genes included RNase III, AGO2, AGO5, DCL1, THO complex protein 3 (THO3) (VIT_19s0015g01220), and the grapevine homolog of Arabidopsis SE (Fig. 6-2). In the Farm 1 samples, two additional upregulated genes, including AGO7 (VIT_03s0038g008300), and six downregulated genes—including RDR2, AGO9 (VIT_06s0009g01200), and a plant combinatorial and modular protein (PCMP-E32,

VIT_07s0130g00180)—were differentially expressed. In the Farm 2 samples, DGE cases included one additional upregulated gene, a ribonuclease (VIT_14s0068g00720) acting as a positive regulator of mRNA decapping, and three downregulated genes, including an uncharacterized protein with 50% similarity to DCL4 of *Triticum urartu* (VIT_11s0149g00100) and two isoforms of RDR1 (VIT_01s0011g05870, VIT_01s0011g05880) (Fig. S5-2, supplementary file).

##### In ‘Vidal’ (V)

15 genes involved in RNAi pathways were significantly regulated (nine upregulated and six downregulated), with 100% of the downregulated genes related to small interfering RNA (siRNA) production and RNA-directed RNA polymerase activity, while 18% of upregulated genes involved grapevine defence responses to viral infection. Significantly (p < 0.01) upregulated genes included DCL1, AGO2, an uncharacterized protein 90% identical to exonuclease-7 (VIT_14s0060g01450), a ribonuclease 3-like protein (VIT_12s0059g01120), RTL2 (VIT_07s0129g00400), and AGO2 (VIT_10s0042g01180). Significantly downregulated genes included AGO9, PCMP-E32, an AGO protein named PNH1 that was 50% identical to AGO10 (VIT_11s0016g04620), two FDM1 isoforms, an uncharacterized RDR-like protein (VIT_11s0016g03220), and an uncharacterized Argo-naute-like protein 50% identical to AGO4 (VIT_08s0040g00070) (Fig. 6-3, supplementary file). In the Farm 1 samples, four additional upregulated genes were identified, including AGO7 and the SICKLE plant protein (VIT_18s0001g07660), along with eight downregulated genes, including DLC2 (VIT_04s0023g00920), RDR1, PCMP-E32, and an uncharacterized double-stranded RNA-binding (DRB) protein (VIT_14s0006g01440) that was 50% identical to *Coffea arabica* DRB4. In the Farm 2 samples, five additional upregulated genes were found, including a ribonuclease (VIT_14s0068g00720) acting as a positive regulator of mRNA decapping and an uncharacterized DRB (VIT_18s0001g07670) that was 50% iden-tical to *Davidia invilucrata* DRB1, along with seven downregulated genes, including RDR6, IND2 (VIT_13s0019g01610), an uncharacterized AGO protein (VIT_06s0061g01040), DCL2, and an uncharacterized retrotransposon-like (RTL) gene (VIT_17s0000g08000) (Fig. S5-3, supplementary file).

#### DGE profile for viruses

##### In ‘Seyval Blanc’

In the Farm 1 samples, three viral genes were significantly (P < 0.01) regulated (one upregulated and two downregulated). The upregulated gene was a GRSPaV coat protein (Rspav1gp5), while the downregulated genes included a GRSPaV helicase pro-tein (Rspav1gp2) and a GSyV helicase protein (GSyV-1_gp1) (Fig. 7-1A-C). In the Farm 2 samples, six viral genes were significantly (P < 0.01) regulated (three upregulated and three downregulated). The upregulated genes included a GSyV helicase protein (GSyV-1_gp1), a polyprotein of grapevine rupestris vein feathering virus (B5P33_gp1), and a GRSPaV coat protein (Rspav1gp5), while the downregulated genes included a GAMaV coat protein (BLM49_gp1), a GRSPaV helicase protein (Rspav1gp2), and a GRSPaV movement protein (Rspav1gp4) (Fig. 7-2 A-C).

**FIG 7.**
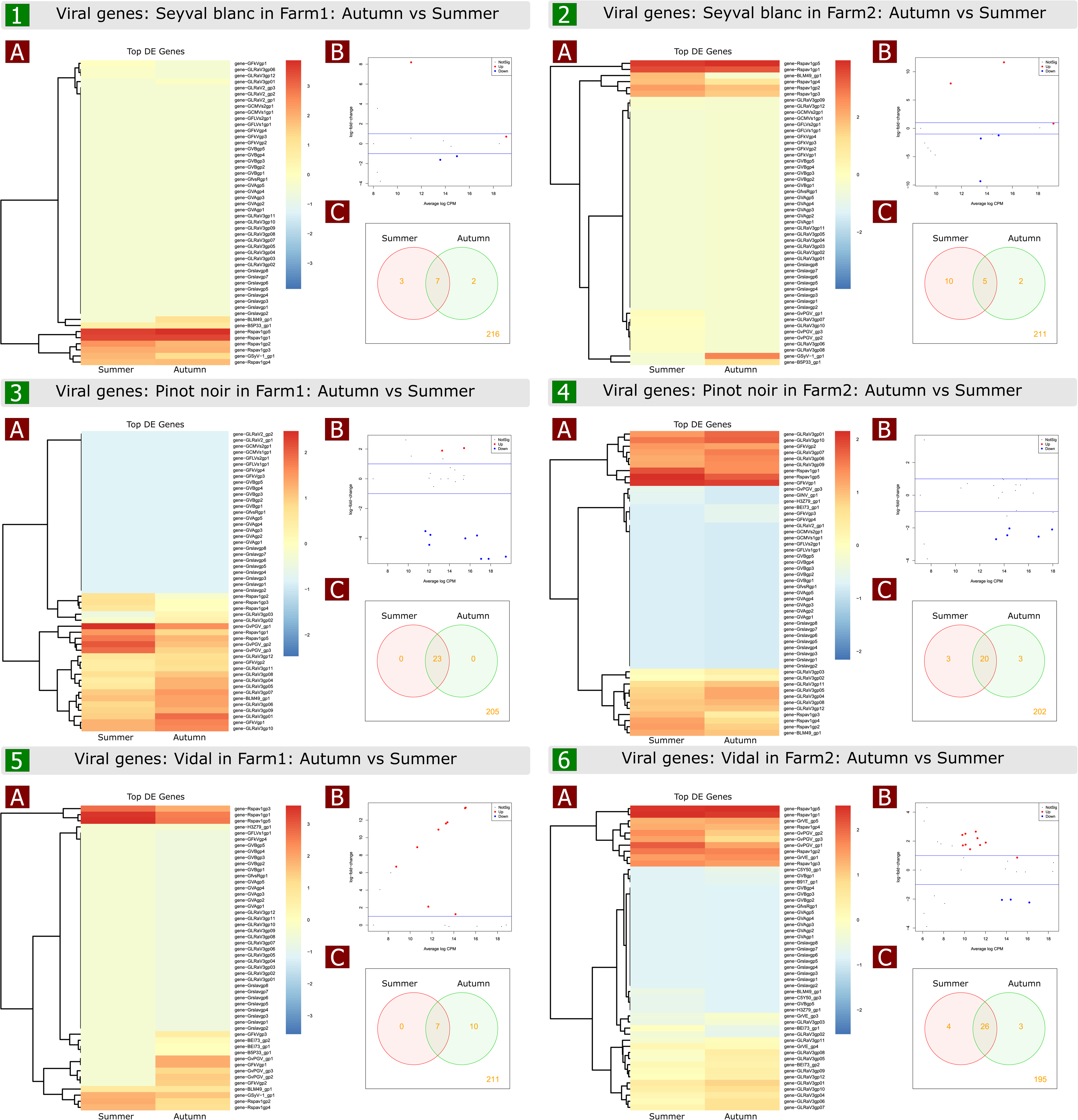
Graphical representations of viral gene groups differentially expressed at the two sampling times (summer vs. autumn) grouped by farm and cultivar. **Panels 1** and **2** represent ‘Seyval Blanc’ (SB) at Farm 1 and 2, respectively. **Panels 3** and **4** represent ‘Pinot Noir’ (PN) at Farm 1 and 2, respectively. **Panels 5** and **6** represent ‘Vidal’ (V) at Farm 1 and 2, respec-tively. **A**, heatmap of the most significant differentially expressed genes in each sample. **B**, mean-difference plot (MD-Plot) for all differentially expressed genes. Red and blue dots in-dicate genes with significant differential expression at the p < 0.05 statistical level. **C,** Venn diagram of number of genes present at each sampling time. The number of genes that are not present in this comparison is shown on the bottom right-hand side.

##### In ‘Pinot Noir’

In the Farm 1 samples, 13 viral genes were significantly (P < 0.01) regulated (five upregulated and eight downregulated). The upregulated genes included GLRaV3 RDR polymerase (GLRaV3gp01), GLRaV3 heat shock protein 70 (GLRaV3gp04), GLRaV3 heat shock protein 90 (GLRaV3gp05), a GLRaV3 coat protein (GLRaV3gp06), and a GLRaV3 small hydrophobic protein (GLRaV3gp03), while the downregulated ones included a GPGV RDR polymerase (GvPGV_gp1), a GPGV movement protein (GvPGV_gp2), a GPGV coat protein (GvPGV_gp3), a GRSPaV coat protein (Rspav1gp5), a GRSPaV replicase (Rspav1gp1), a GRSPaV helicase protein (Rspav1gp2), GRSPaV movement protein TGB2 (Rspav1gp3), and GRSPaV movement protein TGBp3 (Rspav1gp4) (Fig. 7-3 A-C). In the Farm 2 samples, 11 viral genes were significantly (P < 0.01) regulated (three upregulated and eight downregulated). The upregulated genes included a GLRaV3 RDR polymerase (GLRaV3gp01), GLRaV3 heat shock protein 70 (GLRaV3gp04), and GLRaV3 heat shock protein 90 (GLRaV3gp05), while the downregulated ones included a GRSPaV replicase (Rspav1gp1), a GRSPaV helicase protein (Rspav1gp2), GRSPaV movement protein TGB2 (Rspav1gp3), GRSPaV movement protein TGBp3 (Rspav1gp4), a GRSPaV coat protein (Rspav1gp5), a GAMaV coat protein (BLM49_gp1), a GFkV replicase (GFkVgp1), and a GFkV coat protein (GFkVgp2) (Fig. 7-4 A-C).

##### In ‘Vidal’

In the Farm 1 samples, nine viral genes were significantly (P < 0.01) regulated (nine upregulated and none downregulated). The upregulated genes included a GPGV RDR polymerase (GvPGV_gp1), a GPGV movement protein (GvPGV_gp2), a GPGV coat protein (GvPGV_gp3), a GFkV replicase (GFkVgp1), a GFkV coat protein (GFkVgp2), an uncharacterized GFkV ORF3 protein (GFkVgp3), a GAMaV coat protein (BLM49_gp1), a GSyV helicase protein (GSyV-1_gp1), and a grapevine Red Globe virus coat protein (BEI73_gp2) (Fig. 7-5 A-C). In the Farm 2 samples, 13 viral genes were significantly (P < 0.01) regulated (10 upregulated and three downregulated). The upregulated genes included a GLRaV3 RDR polymerase (GLRaV3gp01), GLRaV3 heat shock protein 70 (GLRaV3gp04), GLRaV3 heat shock protein 90 (GLRaV3gp05), a GLRaV3 coat protein (GLRaV3gp06), a diverged copy of GLRaV3 coat protein (GLRaV3gp07), a GLRaV3 p21 protein (GLRaV3gp08), a GLRaV3 p19.6 protein (GLRaV3gp09), a GLRaV3 p19.7 protein (GLRaV3gp10), a GLRaV3 7kDa protein (GLRaV3gp12), and a grapevine virus E RDR polymerase (GrVE_gp1). The down-regulated genes included a GPGV RDR polymerase (GvPGV_gp1), a GPGV movement protein (GvPGV_gp2), and a GPGV coat protein (GvPGV_gp3) (Fig. 7-6 A-C).

Among the viral DGE genes, 12 (viral replicases, viral suppressor of RNA silencing) were validated by quantitative PCR (Fig. 5D and E).

### Metabolome profile

To compare the metabolome profile with the viral and host gene expression profiles, a non-parametric correlation analysis was carried out using viromic data, qPCR results and the metabolomic fold changes detected between the two sampling times (Fig. 8A). The results of the Spearman rank correlation analysis showed that changes in abscisic acid (ABA) concentrations were positively correlated with AGO 10 (0.94, P < 0.001) and SDE3 (0.83) and negatively correlated with HSVd (-0.94, P < 0.001) and THO3 (-0.89, P < 0.001); THO3 was negatively correlated with a GPGV coat protein (GvPGV_GP3) (-0.83, P < 0.001) (Fig. 8A and Fig. S6). Changes in salicylic acid concentrations were negatively correlated with AGO1 (-0.94, P < 0.001). Changes in gallic acid concentrations were negatively correlated with HSVd fold changes (-0.89, P < 0.001), and positively correlated with a GPGV RDR poly-merase (GvPGV_gp1) (0.83, P < 0.001), a GPGV movement protein (GvPGV_gp2) (0.83, P < 0.001), and host RDR2 (0.83, P < 0.001). Luteolin-7-glucoside was negatively correlated with GAMaV fold changes (-0.81) (Fig. 8A, Fig. S6). Unlike luteolin-7-glucoside, luteo-lin-7-glucuronide was negatively correlated (-0.89, P < 0.01) with plant RDR1 (Fig. 8A). Myricetin was positively correlated with fold changes in GLRaV3 (0.82, P < 0.001), a GRSPaV replicase (Rspav1gp1), and a GRSPaV coat protein (Rspav1gp5) (0.94, P < 0.001) and negatively correlated with fold changes in AGO2 and AGO5 (-0.94, P < 0.001). Caffeic acid was negatively correlated with fold changes in GLRaV3 RDR polymerase (GLRaV3gp01) (-0.81, P < 0.001). The total phenol content was negatively correlated with GLRaV3 fold changes (-0.88, P < 0.001), which were negatively correlated with IDN2 fold changes (-0.82, P < 0.001). In contrast, the total phenol content was positively correlated with GPGV fold changes (0.94, P < 0.001) (Fig. 8A, Fig. S6).

**FIG 8.**
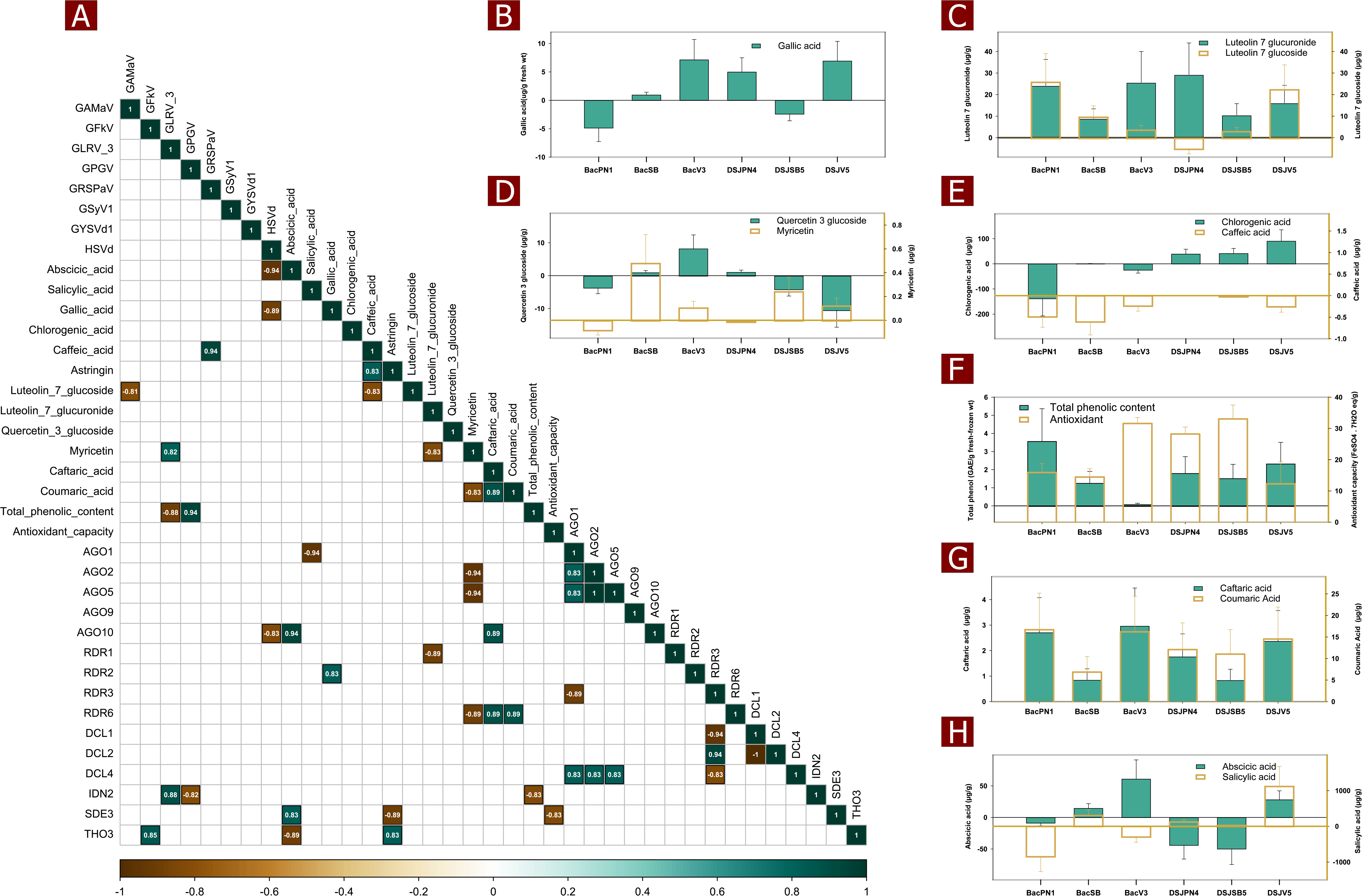
Comparison of metabolite concentrations, viral species titres and host gene expression at the two sampling times. **A**, Spearman rank correlation matrix of viral and host gene ex-pression profiles and metabolites. The colour scale represents the Spearman coefficient of correlation (lower left for r = 1 to upper right for r = −1). Coloured squares represent signifi-cant coefficients at the p < 0.01 statistical level, while blank squares were not significant. **B**, histogram of variations in gallic acid concentrations. **C**, histogram of variations in flavone compound concentrations (luteolin -7-glucuronide and luteolin-7-glucoside). **D**, histogram of variations in flavonol compound (Quercetin 3 glucoside and Myricetin) concentrations. **E**, histogram of variations in organic acid compound (chlorogenic acid and caffeic acid) con-centrations. **F**, histogram of total phenolic content and variations in antioxidant capacity. **G**, histogram of variations in phenolic acid (caftaric acid and coumaric acid) concentrations. **H**, histogram of variations in phytohormone (abscicic acid and salicylic acid) concentrations.

The metabolome profile changed significantly between the two sampling times, depending on the type of metabolic compound considered (Fig.8 B-H). Gallic acid production increased in all plants, except for the PN cultivar at Farm 1 and the SB cultivar at Farm 2 (Fig. 8B). Concentrations of the flavone compounds luteolin-7-glucuronide and luteolin-7-glucoside increased, except in PN at Farm 2, where the production of luteolin-7-glucoside decreased (Fig. 8C). In the case of flavonol compounds, quercetin-3-glucoside concentrations increased in plants BacSB, BacV3 and DSJPN4 and decreased in plants BacPN1, DSJSB5 and DSJV5, while myricetin concentrations decreased only in BacPN1 (Fig. 8D). Among organic acid compounds, the concentrations of chlorogenic acid and caffeic acid decreased in all cultivars at Farm 1, while at Farm 2, concentrations of chlorogenic acid increased and concentrations of caffeic acid decreased or remained unchanged (Fig. 8E). The total phenolic content and the antioxidant capacity increased in all plants, except BacV3, in which the total phenolic acid content at the two sampling times was not significantly different (Fig. 8F). Symptoms were observed in this plant at both sampling times. Phenolic acid production (caftaric acid and coumaric acid) increased in all tested plants (Fig. 8G), which was corroborated in the transcriptomic results. Concentrations of the phytohormone abscisic acid (ABA) decreased in the PN plants, but increased in the V. plants. In the SB plants, ABA concentrations increased in the Farm 1 plants and decreased in the Farm 2 plants. For the phytohormone salicylic acid, concentrations decreased or remained unchanged in all plants, except the SB cultivar at Farm 1 and the V cultivar at Farm 2 (Fig. 8H).

## DISCUSSION

In grapevine, mixed viral infections in the same plant are common ^13^. Our understanding of the factors and signaling pathways driving virus-like symptoms in mixed infections is still sparse and not fully comprehend. In addition, the confounding, antagonistic, or synergistic effects of viruses on symptom expression in mixed infections is not well documented, even for viruses known to cause distinctive symptoms, such as GLRaV3 and GPGV ^14^. In this study, we attempted to determine whether the composition and abundance of the grapevine virome changed over time in mixed infections and whether these changes were associated with symptom expression. We also sought to identify differences in the transcriptomes and metabolomes of asymptomatic and symptomatic plants. Unlike previous studies that compared the transcriptomes of virus-infected and virus-free grapevines, this study compared the transcriptome profiles of the same virus-infected grapevine at two different phenotypic stages (asymptomatic and symptomatic), in order to enhance our understanding of symptom expression in mixed viral infections.

We examined the prevalence of viral infections in grapevine plants at two sampling times (summer and autumn) and evaluated temporal changes in the grapevine virome. In all, 10 viral species and two viroids were detected, including grapevine asteroid mosaic-associated virus, grapevine fleck virus, grapevine leafroll-associated virus, and grapevine Pinot gris virus, with mixed infections being prevalent in most sampled grapevines. The dominance of grapevine rupestris stem pitting-associated virus, grapevine yellow speckle viroid 1, hop stunt viroid, and grapevine leafroll-associated virus 3 in the samples is in line with previous studies, which have found GRSPaV, GYSVd1 and HSVd to be common in grapevine ^4, 15–17^. Re-gardless of the grapevine cultivar, a very diverse virome was detected at both the asymptomatic and symptomatic stages and a large majority (83.33%) of the plants expressed virus-like symptoms in autumn. This is consistent with the findings of several studies ^18, 19^ that observed that virus-like symptoms were more prevalent in grapevine in autumn than in summer. Similarly, other studies ^20, 21^ have observed leafroll symptoms during the fruit ripening, or post-veraison, period, which typically occurs in autumn in northern viticulture. However, our results show that, in 75% of cases, viral abundance either decreased or remained the same between sampling times (summer and autumn), even though 87.5% of the symptomatic plants expressed symptoms in autumn. In addition, no significant associations were found between the increased relative abundance of GLRaV3 and GPGV, which are known to cause symp-toms, and symptom expression. These results are surprising given the findings in the scientific literature that virus titre is correlated with symptom expression and that symptomatic vines contain significantly more virus than asymptomatic vines. Indeed, previous studies demonstrated that grapevines showing symptoms of GLRaV3 and GPGV had high titres of these viruses, while asymptomatic grapevine plants had undetectable virus titres ^22, 23 24^. This result has a major impact on grapevine viruse diagnostics, because the ability to detect viruses using conventional methods (e.g. PCR, ELISA etc.) is hindered by low virus titres. Another potentially important factor that could explain the timing of symptom expression is co-infection with different viruses. Even though most of the viruses and viroids detected in our samples were members of the background virome (HSVd, GYSVd, GRSPaV, etc.) and are not known to cause symptoms in the cultivars used in our study, the impacts of these viruses in mixed infections are unknown. Therefore, we hypothesize that the change in virome composition particularly in the background virome between the two sampling times may explain the discrepancy in the timing of symptoms. However, the different diversity indices used to measure the similarity of the viral communities found at the two sampling times show very similar values for the indices of species richness, composition, and abundance. Consequently, symptom expression in the autumn is not associated with viral titre (even for viruses known to cause symptoms such as GLRaV3 and GPGV) or changes in virome composition. Although we did not evaluate the impact of the intra-host viral haplotype population, especially for GLRaV3 and GPGV, the occurrence of single nucleotide polymorphism in the same viral species over time may explain the timing of symptoms, as we recently demonstrated for GPGV ^25^. Furthermore, symptom expression may also be a consequence of leaf malfunction and metabolic compound accumulation in response to viral infections, as shown in the recent model proposed by Song et al. ^19^.

The analysis of differential gene expression in grapevines shows that cell wall biosynthesis (e.g. hemicellulose, polysaccharide, chitin, pectin) pathways were significantly downregulated in all GRLaV3, GAMaV or GPGV-infected grapevine plants, while these pathways were significantly upregulated in plants free of these viruses (SB, Farm 1). These results are in line with previous studies that found that the genes involved in cell-wall softening and expansion in GLRaV3-infected grapevine leaves were downregulated, suggesting that the presence of other symptomatic viruses (e.g. GAMaV and GPGV) was not interfering with GLRaV3 symptomatology ^14, 19^. In addition, polyketide (e.g. flavonoid, stilbenoid, and gingerol biosynthesis) and carbohydrate catabolic pathways were significantly upregulated in all cultivars. The polyketide pathway was upregulated in SB, V and Pn by 12%, 21%, and 23%, respectively. The SB plants were GLRaV3- and GPGV-free. These results suggest that the upregulation of polyketide biosynthesis is more pronounced in GLRaV3- and GPGV-infected grapevines, which is consistent with other studies that have found an upregulated polyketide pathway in GLRaV3-infected grapevine ^26, 27^. In dark-skinned cultivars, numerous studies have shown that the upregulation of flavonoid biogenesis seems to trigger red-purple discoloration; however, this phenomenon has not been observed in GRLaV3-infected hybrid white cultivars ^19, 28^. Furthermore, total phenolic content and antioxidant capacity increased in all plants, except for BacV3, which, unlike the other plants, displayed early-onset symptoms (summer), suggesting the potential role of phenolic content in virus-like symptom expression. Indeed, total phenolic content values were negatively correlated with GLRaV3 fold changes and positively correlated with GPGV fold changes. This result contrasts with the recent findings of Wallis (2022), who observed reduced total phenolic levels in virus-infected grapevines. In addition, the production of flavonoid precursors (e.g. abscisic acid, or ABA) was also significantly reduced in plants expressing strong virus-like symptoms (BacPN1, DSJPN4, DSJSB5), in conjunction with reduced yield (cluster size, density, and number), except for plant BacV3, which displayed GPGV symptoms (no yield reduction) despite a significant increase in ABA concentrations. However, as mentioned above, BacV3 exhibited symptoms at both sampling times. This result is line with the conclusion of a recent study on GLRaV-grapevine interactions in a red cultivar ^20^, which supports the role of ABA in the shared responses to GLRaV infection. Indeed, ABA can interact with RNA-silencing mechanisms (RNAi) and trigger antiviral defence responses ^20, 29^. Consequently, transcripts involved in the RNAi mechanism are likely significantly regulated.

A total of 31 (out of 65) genes involved in RNA-silencing mechanisms were differentially expressed by all the plants. In ‘Seyval Blanc,’ 14 RNAi-associated genes were significantly regulated, with 42% of the downregulated genes related to grapevine defence responses to viral infection. In ‘Pinot Noir,’ 80% of downregulated genes were related to these defence responses, while only 18% of the genes in ‘Vidal’ related to defence responses were upregulated. In the ‘Vidal’ plant at Farm 1 (BacV3), which displayed strong GPGV symptoms in both summer and autumn, nine and zero viral RNAi-related genes were significantly upregulated and downregulated, respectively. The upregulated viral genes included GPGV RDR polymerase and the GPGV movement protein and coat protein (GvPGV_gp3) genes; the coat protein gene has recently been proven, based on in silico prediction, to suppress RNA silencing in *Nicotiana binthamiana* plants ^30^. In contrast, the GPGV- and GLRaV3-infected ‘Vidal’ at Farm 2 (plant DSJV5) expressed mild symptoms only in autumn, with all GPGV genes, including RDR polymerase and movement and coat proteins, downregulated. The only difference in the transcriptional profile of the two plants (DSJV5 and BacV3) was the upregulation of an uncharacterized double-stranded RNA-binding protein in DSJV5, which was downregulated in BacV3. In fact, this protein is required to assist Dicer-like protein 4 (DLC4) in directing the selection and loading of dsRNA to the Argonaute protein for the degradation and generation of small interfering RNAs against viral infection in *Arabidopsis thaliana* ^31–33^. However, to our knowledge, the putative role of this double-stranded RNA-binding protein in grapevine has never been elucidated despite its documented homologs in the grapevine genome. This result also reinforces the recent hypothesis that dou-ble-stranded RNA-binding proteins play a critical role in establishing dominant antiviral responses in plants ^34^. Overall, these results suggest that the activation of host antiviral defence responses, viral counter-defence responses, and symptom development are concomitant.

The non-parametric correlation analysis revealed various correlations between changes in the concentration of different metabolites and fold changes in viral and host gene expression. For instance, changes in abscisic acid concentrations were positively correlated with the AGO10 and SDE3 genes, and negatively correlated with HSVd and the THO3 gene, which, in turn, were negatively correlated with the GPGV coat protein. On the other hand, changes in salicylic acid concentrations were negatively correlated with the AGO1 gene, while changes in gallic acid concentrations were negatively correlated with HSVd and positively correlated with GPGV RDR polymerase, a GPGV movement protein, and host RDR2. Luteo-lin-7-glucoside was negatively correlated with GAMaV fold changes, while myricetin was positively correlated with GLRaV3 fold changes, and GRSPaV replicase and coat protein fold changes. These findings are consistent with previous studies and support the hypothesis that antiviral responses (RNAi) in grapevines induce oxidative stress leading to symptomatic leaves ^21, 35, 36^. However, this hypothesis has not been verified in any published studies in a mixed viral context, even though individual virus-host interactions are known to lead to distinct outcomes.

In conclusion, these results provide insights into the complex gene expression patterns in grapevines and highlight the importance of considering cultivar and farm effects when studying virus-host interactions. Unlike the majority of studies on grapevine-virus interactions, which are conducted using samples collected at a single point in time, this study involved sampling at two different times of year in order to compare the transcriptome profile. Therefore, we were able to compare not only the transcriptome profiles of different plants at the same phenological stage but also the transcriptome profiles of the same plant at different phenological stages. Our results support the hypothesis that virus-host interactions are modulated in a specific phenological manner ^21^. Therefore, the expression of disease symp-toms in grapevine in autumn is the result of a dynamic interplay between the virome and antiviral factors in the plant that occur at distinct phenological stages.

## MATERIALS AND METHODS

### Plant material

The grapevine plants used in this study were all grown in two different vineyards located in Quebec, Canada. Mature leaf materials (including blades and petioles) from 18 grapevine plants of three different varieties (‘Pinot Noir,’ ‘Vidal,’ and ‘Seyval Blanc’ cultivars) were collected at two times of the year in 2019, for a total of 36 observations. The first set of samples were collected on August 2 (summer) and the second, on September 30 (autumn). Five leaves per plant (three from the bottom, one from the middle and one from the top of the canopy) were harvested and immediately frozen in liquid nitrogen and stored at -80°C for further analysis. All plants were visually inspected for virus-like symptoms, mainly those caused by grapevine leafroll-associated viruses and GPGV.

### Nucleic acid extraction, library preparation and sequencing

#### Double-stranded RNA (dsRNA) extraction

Grapevine leaves were individually washed with distilled water and crushed before being homogenized in a liquid-nitrogen-cooled 50-mL stainless-steel grinding jar, following the protocol described in Fall et al. ^4^. The dsRNA was then extracted using a modified version of the technique developed by Kesanakurti et al. ^37^ as described in Fall et al. ^4^. Briefly, black bean (*Phaseolus vulgaris*) was added as a positive control to the homogenized sample-extraction buffer mix at a final concentration of 1% (w/w). The endornavirus species *Phaseolus vulgaris endornavirus* 1 (PvEV1), an endophyte virus of *P. vulgaris*, was used to assess the effectiveness of dsRNA extraction and to monitor the accuracy of the high-throughput sequencing results.

#### Total RNA extraction

Total RNA was extracted from a total of six randomly selected grapevine plants expressing characteristic virus-like symptoms at different levels of severity. For all the samples, ∼50 mg of tissue was ground under liquid nitrogen and the total RNA was extracted using the Spectrum Plant Total RNA Kit (Sigma-Aldrich) following the manufacturer’s protocol. RNA quality and quantity were determined using a NanoDrop 2000 Spectrophotometer, 1% agarose gel, and a Bioanalyzer Chip RNA 7500 Series II (Agilent). A threshold RNA integrity number (RIN) ≥ 5 (Bioanalyzer) was used to assess RNA integrity and suitability for RNASeq library preparation.

#### Library preparation and sequencing

The dsRNA was denatured for cDNA synthesis and treated with RNase H following the protocol described in Fall et al. ^4^. Libraries were constructed using the Nextera XT DNA Library Preparation Kit (Illumina) with 1 ng of input double-stranded cDNA, and sequencing was performed using MiSeq 2 × 250 cycle paired-end (Illumina) with the Nano Kits v3 reagent. The black bean added to all grapevine samples was sequenced separately to differentiate its virome from the grapevine virome associated with each sample ^4^. The total RNA extracted was sent to Genome Quebec for library synthesis and RNA sequencing using the Illumina NovaSeq platform.

### Bioinformatics and statistical analysis

A number of packages using R software, version 4.0.3, were utilized to perform the statistical tests and generate the graphics used in this study, notably ggplot2, vegan, gplots, RcolorBrewer, and ape.

#### Processing of raw data

All the raw data (FASTQ) files were demultiplexed using FastQC (v0.11.8) (ref: https://www.bioinformatics.babraham.ac.uk/projects/fastqc/). FastQC was also used to evaluate the quality of the reads. Subsequently, read quality trimming (minimum score of 30) was carried out with Trimmomatic V.0.32. Next, AfterQC (v0.9.7) (ref: https://doi.org/10.1186/s12859-017-1469-3) was used to remove and trim the low-quality sequences and read adapters.

##### Grapevine virome characterization and statistical analysis

The paired FASTQ files resulting from dsRNA sequencing that passed quality control were analyzed using two different pipelines ^38, 39^ as described in Fall et al. ^4^. The criteria for positive detection of a virus were detection by both pipelines with a whole-genome coverage of at least 0.5 and a weight (titre) greater than 0.001, as well as inclusion in a sample positive for PvEV1 (the positive control) ^4, 39^. Depth, or the number of times a genome is covered by mapped reads, was used as a measure of absolute abundance and, since many multivariate methods are sensitive to total abundance (Kembel and Cahill 2011), the relative abundance of each virus detected in each sample was calculated using the corresponding function in the vegan package. After relative abundance was normalized using the “order” function in the vegan package, ggplot2 functions were used to generate graphs of the relative abundance (proportions) of viral species as a function of the sample. To compare viral abundance at the two different times of the year that samples were taken (summer and autumn), a non-parametric test (Kruskal-Wallis test) was performed iteratively using the kruskal.test function, with the value of alpha set at 0.01. The p-value, the number of false positives expected by chance (E-value) and the probability of finding at least one false positive by chance were calculated (family-wise error rate [FWER]). A significant difference in viral abundance in summer and fall is indicated by an E-value much smaller than 1; the q-value, which is the adjusted p-value after the Benjamini and Hochberg correction, was also considered ^40, 41^. In addition, non-metric multi-dimensional scaling (NMDS) was used to plot dissimilarities in viral abundance. A Bray-Curtis dissimilarity matrix was generated and the covariance matrix was calculated using the corresponding function in the vegan package ^42, 43^.

Three different community diversity indices were calculated to evaluate how the virome evolved between the two times in the year that vines were sampled. First, we calculated the Gini-Simpson index, a measure of the probability that two randomly selected individuals will belong to the same species ^44^:

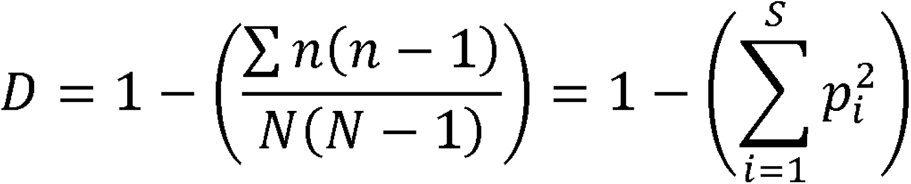

Where *D* is the Gini-Simpson index, *n* is the total number of specimens of a given virus species, *N* is the total number of specimens of all virus species, *s* is the number of species, *pi* is the relative abundance of i-th species, and *i* is the number of individuals of species i. The values for *D* range from 0 to 1 and increase with viral species richness. Secondly, we calculated the Shannon-Wiener diversity index, a measure of the uncertainty associated with the prediction of the identity of a species obtained in sampling; it is less sensitive to the abundance of a given virus species ^44^:

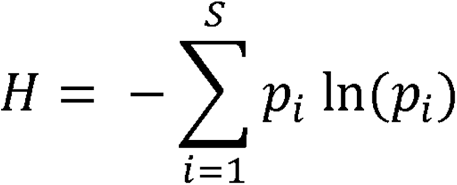

Where *H* is the Shannon-Wiener diversity index, *s* is species richness, and *pi* is the relative abundance of species *i*. The values for *H* range from 0 to 5; *H* = 0 when the virome only contains one species and increases as the community contains more species. Lastly, we used Morista’s index of similarity to measure the similarity of the viral communities (species richness and abundance) at the two sampling times. The value for the Morista-Horn index ranges from 0 (no similarity) to 1 (complete similarity) and is calculated as follows:

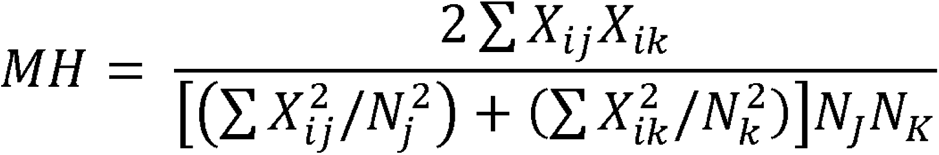

Where *MH* is the Morista-Horn index, *x_ij_* and *x_ik_* are the number of individuals of species i in sample j and sample k respectively, and *N_J_* and *N_K_* are the total number of individuals in samples j and k respectively ^45^.

##### Transcriptome of grapevine virus-infected leaves and statistical analysis

The trimmed reads were mapped to the *Vitis vinifera* PN40024 genome (12x.V2) using the Spliced Transcripts Alignment to a Reference (STAR) aligner (v2.7.9a) ^46^. The resulting BAM files were analyzed using FeatureCounts (v2.0.1) ^47^ and raw counts were obtained for each gene. An EdgeR (v3.34.0) function ^48^ was then used to moderate the degree of overdispersion across transcribed genes, improve the reliability of inference, and complete the downstream analyses in R. Differential gene expression (DGE) analysis was performed in three different sets for a total of 24 separate analyses. These DGE analyses included (i) a comparison of cultivars by farm and sampling time (Fig. S1A); (ii) a comparison of farms at the two sampling times, summer and autumn (Fig. S1B); and (iii) a comparison of sampling times, first, considering all samples; second, by individual cultivar; and, lastly, a pairwise comparison (Fig. S1C). Significant DGEs were sorted, based first on the P-value (0.01) and then using a fold-change threshold of 4. To analyze the DGEs, BiomaRt (v2.48.3) ^49^ was used to retrieve the metadata on gene annotations. The results were visualized using Glimma (v2.2.0) ^50^ and the R package Pheatmap was used to generate heatmaps. Lastly, ClueGO (v2.0.6) (Bindea et al. 2009) was applied to the results of this analysis to examine gene ontology and different aspects of this ontology (including molecular functions, biological processes, and cellular components) were evaluated using cluster analysis. ClueGO was also used for the network analysis of gene ontology terms and pathways in order to investigate DGE in genes involved in RNA interference (RNAi) silencing pathways. A total of 65 different genes involved in these pathways were identified (see supplementary file); an entire set of DGE analyses (see above) were performed on each of these genes. These analyses were also performed on the viruses present in the samples to investigate DGEs.

##### Quantitative RT-PCR of selective DGEs

Among these 65 DGEs, 12 viral genes (viral replicases, viral RNA silencing suppressors) and 22 grapevines genes (Dicer-like proteins, Argonaute proteins, RNA-dependent RNA polymerase) were selected to verify the RNA se-quencing results (supplementary file). The primers used to amplify the targeted genes are listed in the supplementary file. The reference genes VvAIG1 (AvrRpt2-induced gene) and VvTCPB (T-complex 1 subunit beta) were chosen for their highly stable expression in grapevine tissues ^51^. The other reference genes—UBC (ubiquitin conjugating enzyme), VAG (vacuolar ATPase subunit G), and PEP (phosphoenolpyruvate carboxylase)—were selected for their highly stable expression in grapevine when subjected to biotic and abiotic stresses, as well as their previous use as reference genes in other grapevine studies ^52^. The reactions were completed in a final volume of 10 µl using the Luna Universal One-Step RT-qPCR Kit (New England Biolabs), and relative gene expression analysis using the Livak method was performed ^53^. Control gene experiments were also conducted on all samples to ensure the accuracy and reliability of the results.

#### Metabolomic analysis

##### Extraction of grapevine compounds and estimation of antioxidant capacity

For each of the samples phenolic compounds were extracted from 2 g of grape leaves initially crushed in liquid nitrogen and stored at -80 °C according to the protocol adapted from Arigo et al. 2018. The extracts obtained were filtered with a 0.20-µm filter and then analyzed with an Agilent high performance liquid chromatography (HPLC) system equipped with a 150 × 2.1 mm Eclipse Plus phenyl-hexyl column. The mobile phase consisted of solvents A and B (0.1% formic acid in water and 0.1% formic acid in acetonitrile, respectively) and the flow rate was 1.0 mL/min with an injection volume of 20 μL. The gradient program was optimized as follows: linear gradient from 10% to 22% B in 3 min, from 22% to 30% B in 12 min, and from 30% to 90% B in 8 min; 90% isocratic B in 3 min; and linear gradient from 90% to 10% B in 2 min, with a 5-min post-analysis time to equilibrate the system. The diode array detector collected data from 190 nm to 400 nm, while signals were collected at 280 nm and 330 nm for each extract.

Ferric-reducing antioxidant capacity and total phenolic content assays were performed on each sample based on the procedure described by Xie et al. ^54^ with slight modifications. For both assays, 5 g from each frozen sample of grapevine leaves was homogenized with 50 ml of 50% MeOH and then centrifuged at 8,500 rpm at 4 °C for 20 min. After centrifugation, the supernatants were placed in 1.5-mL Eppendorf tubes and stored at −80 °C for the fer-ric-reducing antioxidant capacity and total phenolic content analyses.

To compare the metabolome profile with the viral and host gene expression profile, a correlation analysis was performed, using the Spearman rank correlation method because of its data distribution independence and robustness to outliers. The rcorr and corrplot functions in the Hmisc and corrplot packages were used to generate and visualize the correlation matrix and associated p-value (0.01) and determine the statistical significance of the correlations.

## Supporting information

SupFigure 1

SupFigure 2

SupFigure 3

SupFigure 4

SupFigure 5

SupFigure 6

## Supplementary Materials

The following are available online: Fig. S1, Fig. S2, Fig. S3, Fig. S4, Fig. S5, Fig. S6 and supplementary files containing all relevant data.

## Author Contributions

Conceptualization, XX and Y.Y.; methodology, X.X.; software, X.X.; validation, X.X., Y.Y. and Z.Z.; formal analysis, X.X.; investigation, X.X.; resources, X.X.; data curation, X.X.; writing—original draft preparation, X.X.; writing—review and editing, X.X.; visualization, X.X.; supervision, X.X.; project administration, X.X.; funding acquisition, Y.Y. All authors have read and agreed to the published version of the manuscript.

## Acknowledgments

The authors gratefully acknowledge Joel Lafond-Lapalme, Biology Study Leader in Bioinformatics at Agriculture and Agri-Food Canada, for his help and consistent assistance in bioinformatics data processing. We are also grateful to Eric Courchesne and his team for their support and management of the experimental vineyard. We would also like to express our sincere gratitude to Quebec grapevine growers, the Conseil des vins du Québec (CVQ), and Louis Thomas (chair, CVQ R&D committee), for their support and permission to access their vineyards during a critical period of the production season. We would also like to acknowledge the administrative support that we received from the team at the Saint-Jean-sur-Richelieu Research and Development Centre (the teams led by Vicky Toussaint and Mélanie Cadieux in particular).

## Other information

The raw sequencing data associated with this study has been deposited at the Sequence Read Archive (SRA): Bioproject PRJNA853579, biosamples from SAMN29398272 to SAMN29398307 (SRA from 29398272 to 29398307). Additional data are available in the supplementary file. For any question, please contact the corresponding author.

---

**FIG S1.** Comparison of differential gene expression by cultivar, farm and sampling time. Differential gene expression analysis was performed in three different sets for a total of 24 separate analyses. **A**, comparison of cultivars by farm and sampling time. **B**, comparison of farms at two sampling times (summer and autumn). **C**, comparison of sampling times (1) considering all samples; (2) at the cultivar level; and (3) by pairwise comparison. Significant differential gene expression was determined first by the P-value (0.01) and then a fold-change threshold of 4.

**FIG S2.** Symptomatology of grapevine in leaves collected at Farm 2 at two sampling times. **A** represents plant BacSB (‘Seyval Blanc’), **B** represents plant BacV3 (‘Vidal’) and **C** represents plant BacPN1 (‘Pinot Noir’), samples of which were collected during the summer. **D** represents plant DSJSB5 (‘Seyval Blanc’), **E** represents plant DSJV5 (‘Vidal’) and **F** represents plant BacPN4 (‘Pinot Noir’), samples of which were collected during the autumn. The virome, transcriptome and metabolome profiles of these grapevine plants were determined.

**FIG S3.** Graphical representations of all functional gene groups and pathways differentially expressed at the two sampling times (summer vs. autumn) without consideration of farm location. **Panel 1** represents all cultivars together, **Panel 2** represents ‘Seyval Blanc’ (SB), **Panel 3** represents ‘Pinot Noir’ (PN), and **Panel 4** represents ‘Vidal’ (V). **A**, overview chart with functional groups including specific terms for differentially expressed genes. **B**, gene ontology (GO)/pathway terms (biological process) relating to differentially expressed genes. The bars represent the percentage of differentially expressed genes associated with the terms. The number of differentially expressed genes per term is shown on the bar label. Single (*) or double (**) asterisks indicate significantly enriched GO terms at the p < 0.05 and p < 0.01 statistical levels, respectively. The top and bottom charts show upregulated and downregulated genes, respectively. **C**, graphical representation of interactions among the most significant differentially expressed genes (upregulated and downregulated) in biological processes. GO terms are represented as nodes, and the node size represents the enrichment significance of the term. Only the most significant term in the group was represented. **D**, number of upregulated and downregulated genes in the comparison. **E**, heatmap of the most significant differentially expressed genes in each sample. **F**, mean-difference plot (MD-Plot) for all differentially expressed genes. Red and blue dots indicate genes with significant differential expression at the p < 0.05 statistical level. **G**, Venn diagram of number of genes present at each sampling time. The number of genes that are not present in this comparison is shown on the bottom right-hand side. **H**, Venn diagram of number of genes present at each sampling time (S, summer, A, autumn) in Farm 1 (F1) and Farm 2 (F2). The number of genes that are not present in this comparison is shown on the bottom right-hand side.

**FIG S4.** Graphical representation of all functional gene groups and pathways differentially expressed, grouped by cultivar and farm location. **Panel 1** represents ‘Pinot Noir’ (PN) at Farm 1. **Panel 2** represents ‘Seyval Blanc’ (SB) at Farm 1. **Panel 4** represent ‘Vidal’ (V) at Farm 1. **Panel 4** represents ‘Pinot Noir’ (PN) at Farm 2. **Panel 5** represents ‘Seyval Blanc’ (SB) at Farm 2. **Panel 6** represent ‘Vidal’ (V) at Farm 2. **A**, overview chart with functional groups including specific terms for differentially expressed genes. **B**, gene ontology (GO)/pathway terms (biological processes) relating to differentially expressed genes. The bars represent the percentage of differentially expressed genes associated with the terms. The number of differentially expressed genes per term is shown on the bar label. Single (*) or double (**) asterisks indicate significantly enriched GO terms at the p < 0.05 and p < 0.01 statistical levels, respectively. The top and bottom charts show upregulated and downregulated genes, respectively. **C**, graphical representation of interactions among the most significant differentially expressed genes (upregulated and downregulated) in biological processes. GO terms are represented as nodes, and the node size represents the term’s enrichment significance. Only the most significant term in the group was represented. **D**, number of upregulated and downregulated genes in the comparison. **E**, heatmap of the most significant differentially expressed genes in each sample. **F**, mean-difference plot (MD-Plot) for all differentially expressed genes. Red and blue dots indicate genes with significant differential expression at the p < 0.05 statistical level. **G,** Venn diagram of number of genes present at each sampling time. The number of genes that are not present in this comparison is shown on the bottom right-hand side.

**FIG S5.** Graphical representations of functional gene groups related to the grapevine RNA-silencing pathways differentially expressed at the two sampling times (summer vs. autumn), grouped by cultivar and farm location. **Panel 1** represents ‘Seyval Blanc’ at Farm 1, **Panel 2** represents ‘Pinot Noir’ at Farm 1 and **Panel 3** represents ‘Vidal’ at Farm 1. **Panel 4** represents ‘Seyval Blanc’ at Farm 2, **Panel 5** represents ‘Pinot Noir’ at Farm 2 and **Panel 6** represents ‘Vidal’ at Farm 2. **A**, overview chart with functional groups including specific terms for differentially expressed genes. **B**, gene ontology (GO)/pathway terms (biological processes) relating to differentially expressed genes. The bars represent the percentage of differentially expressed genes associated with the terms. The number of differentially expressed genes per term is shown on the bar label. Single (*) or double (**) asterisks indicate significantly enriched GO terms at the p < 0.05 and p < 0.01 statistical levels, respectively. The top and bottom charts show upregulated and downregulated genes, respectively. **C**, graphical representation of interactions among the most significant differentially expressed genes (upregulated and downregulated) in biological processes. GO terms are represented as nodes, and the node size represents the term’s enrichment significance. Only the most significant term in the group was represented. **D**, number of upregulated and downregulated genes in this comparison. **E**, heatmap of the most significant differentially expressed genes in each sample. **F**, mean-difference plot (MD-Plot) for all differentially expressed genes. Red and blue dots indicate genes with significant differential expression at the p < 0.05 statistical level. **G**, Venn diagram of number of genes present at each sampling time. The number of genes that are not present in this comparison is shown on the bottom right-hand side.

**FIG S6.** Comparison of metabolite concentrations, and the differential expression of viral and host genes at the two sampling times. A Spearman rank correlation matrix was used. The colour scale represents the Spearman coefficient of correlation (lower left for r = 1 to upper right for r = −1). Coloured squares represent significant coefficients at the p < 0.01 statistical level, while blank squares were not significant.

