## Supplementary figures and images for "From Asymptomatic to Symptomatic: Multiomics profiling of the temporal response of grapevine viral-mixed infection"

### SupFigure 1

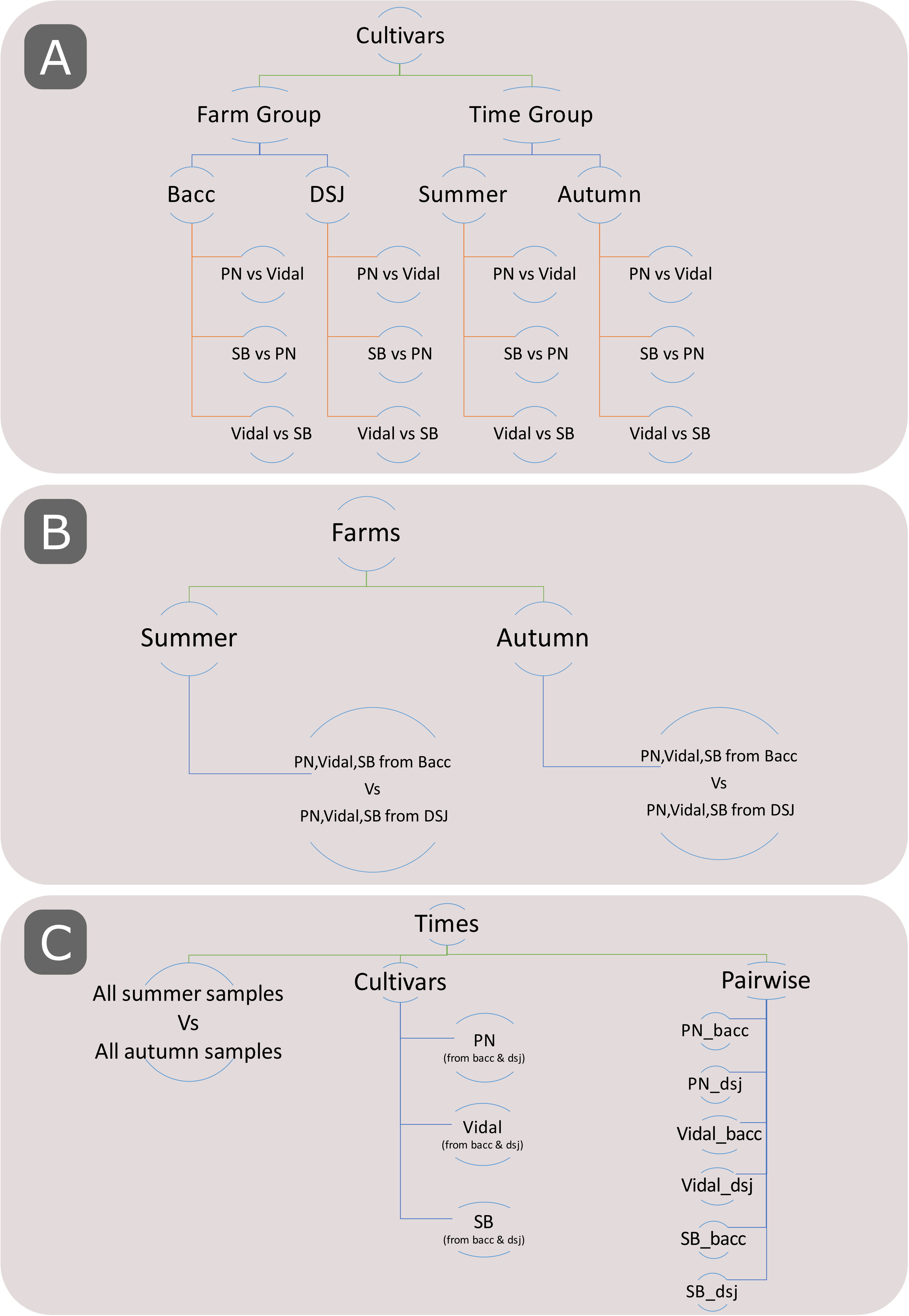

### SupFigure 2

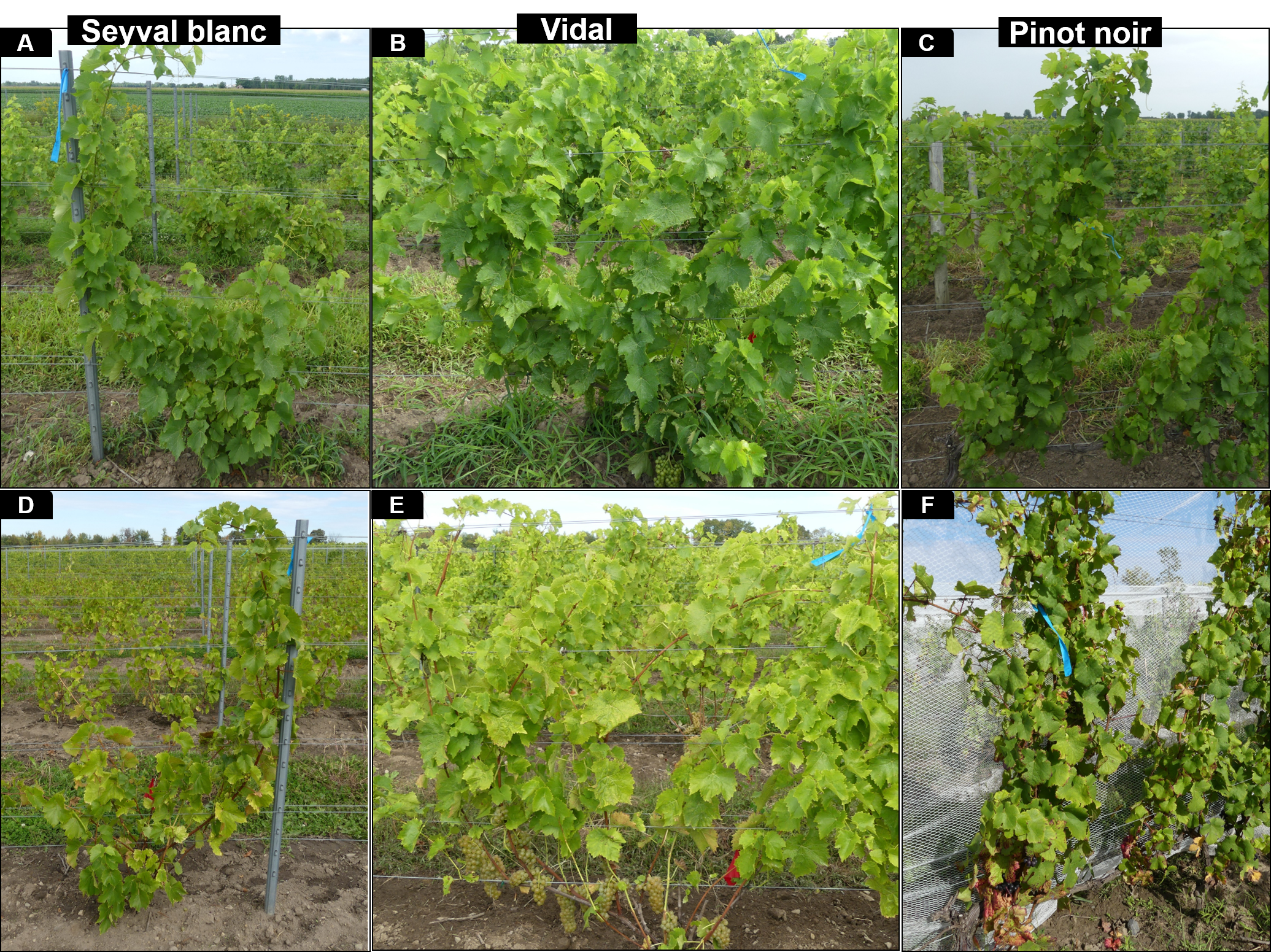

### SupFigure 3

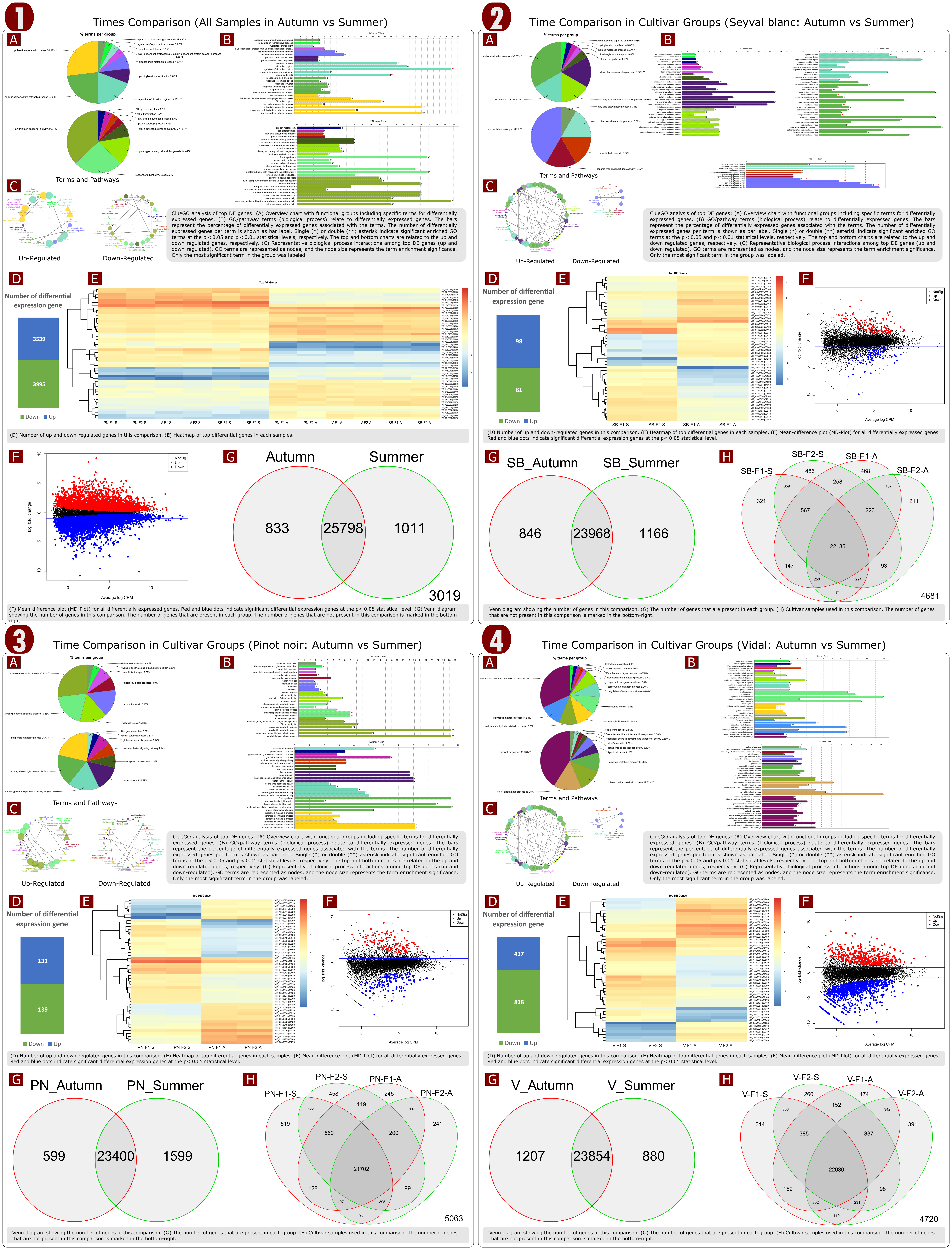

### SupFigure 5

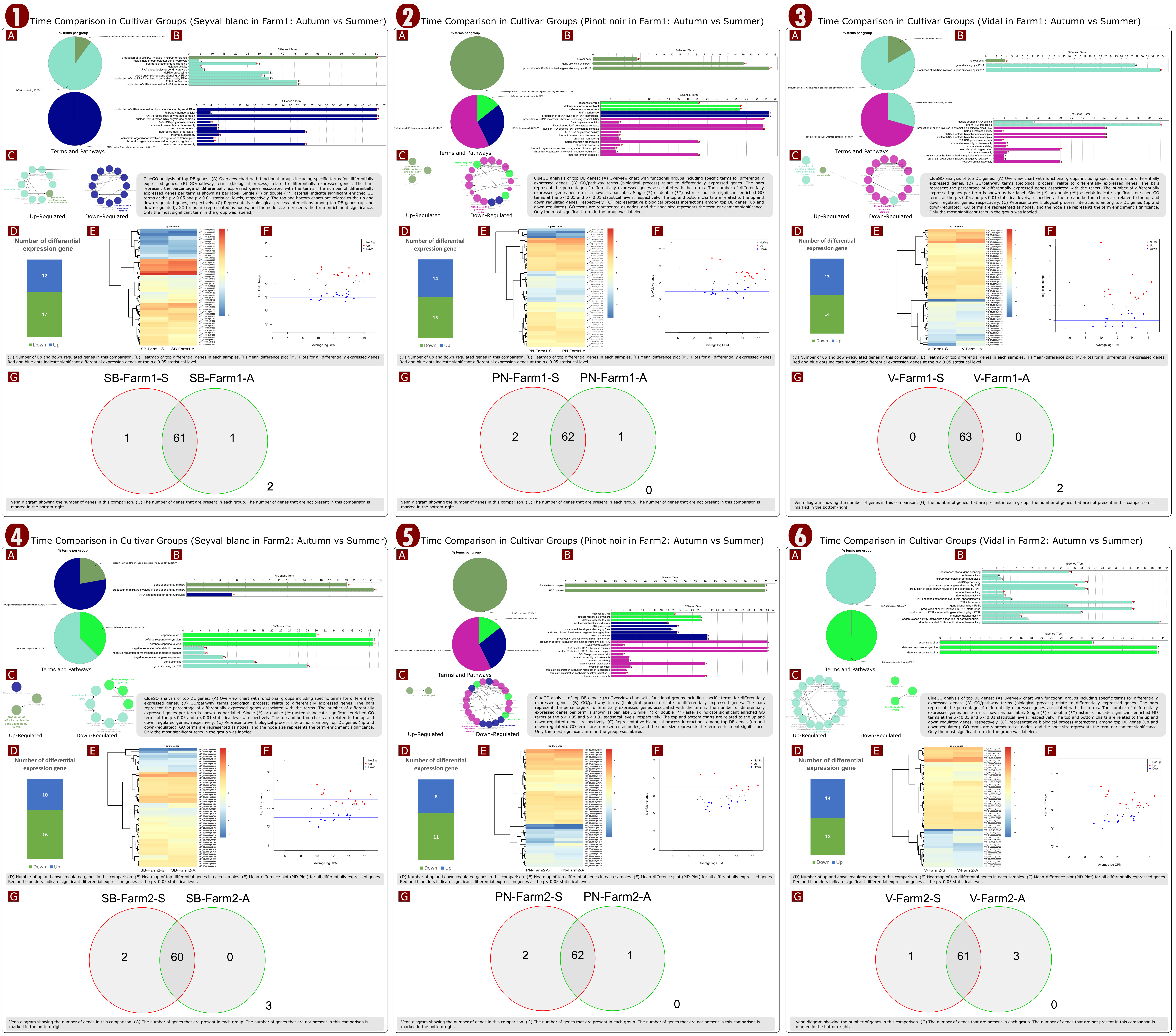

### SupFigure 6

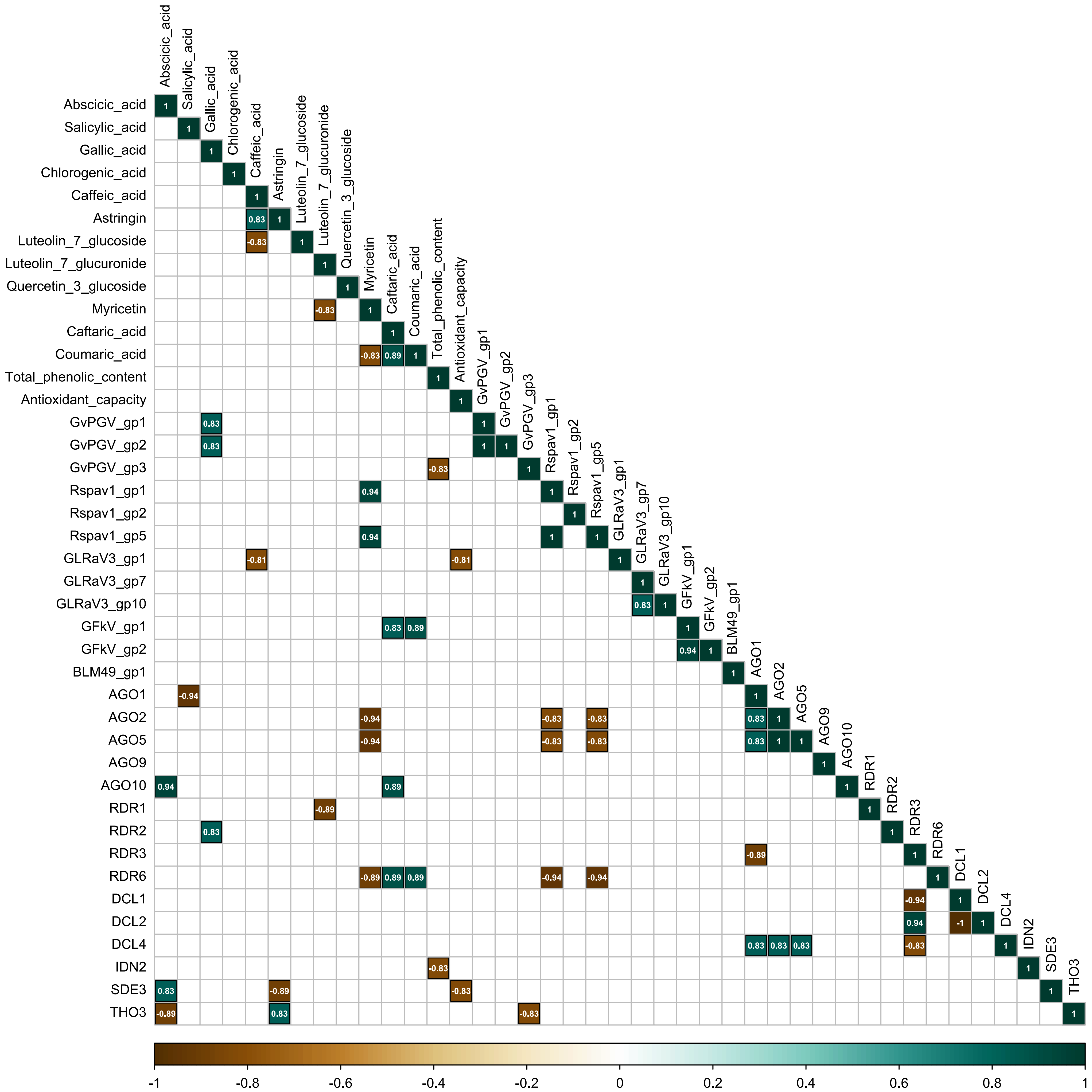
