## Supplementary material for "From Asymptomatic to Symptomatic: Multiomics profiling of the temporal response of grapevine viral-mixed infection": SupFigure 4

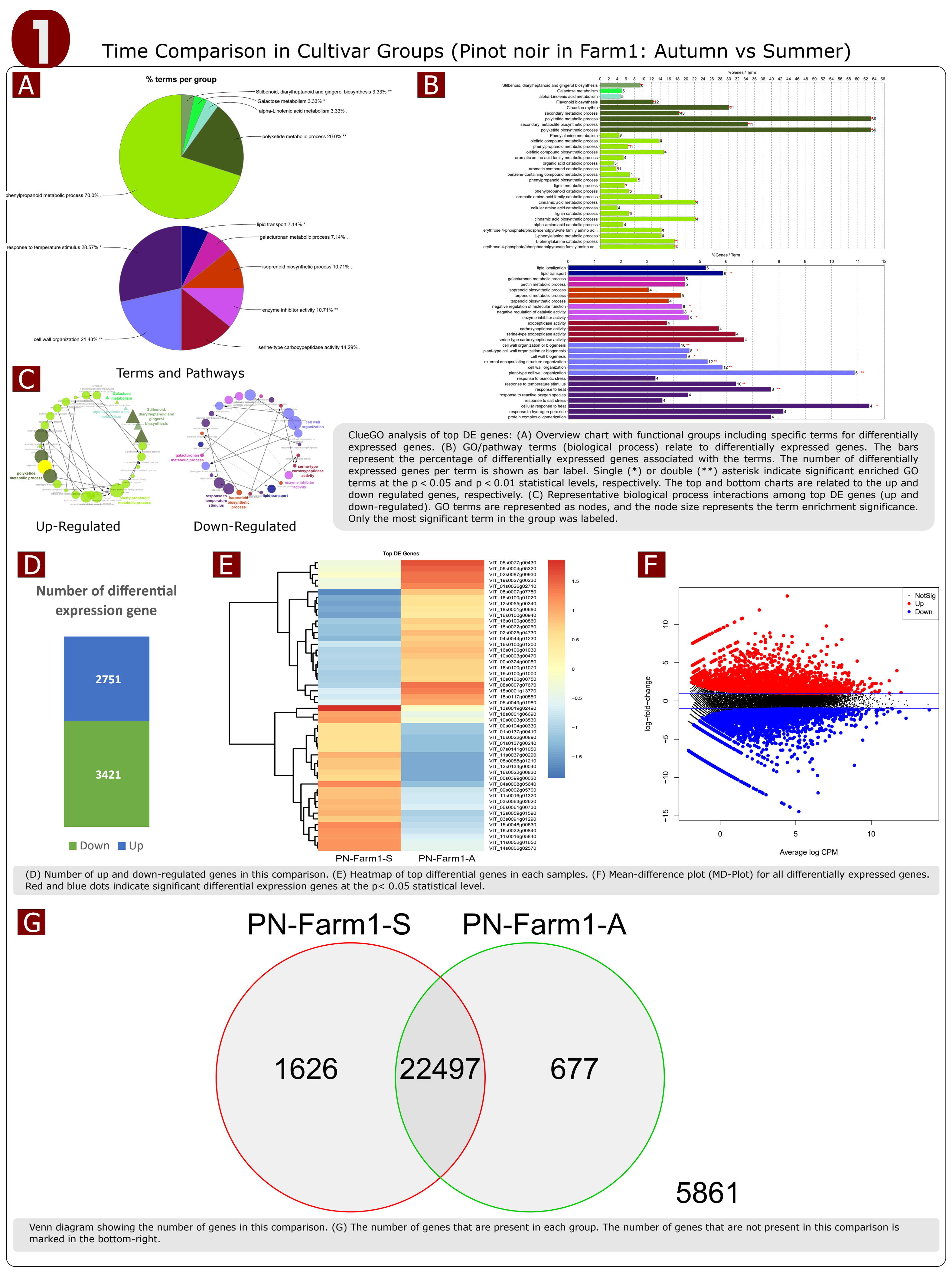


ClueGO analysis of top DE genes: (A) Overview chart with functional groups including specific terms for differentially expressed genes. (B) GO/pathway terms (biological process) relate to differentially expressed genes. The bars represent the percentage of differentially expressed genes associated with the terms. The number of differentially expressed genes per term is shown as bar label. Single (*) or double (**) asterisk indicate significant enriched GO terms at the p < 0.05 and p < 0.01 statistical levels, respectively. The top and bottom charts are related to the up and down regulated genes, respectively. (C) Representative biological process interactions among top DE genes (up and down-regulated). GO terms are represented as nodes, and the node size represents the term enrichment significance. Only the most significant term in the group was labeled.

(D) Number of up and down-regulated genes in this comparison. (E) Heatmap of top differential genes in each samples. (F) Mean-difference plot (MD-Plot) for all differentially expressed genes. Red and blue dots indicate significant differential expression genes at the p< 0.05 statistical level.

Venn diagram showing the number of genes in this comparison. (G) The number of genes that are present in each group. The number of genes that are not present in this comparison is marked in the bottom-right.

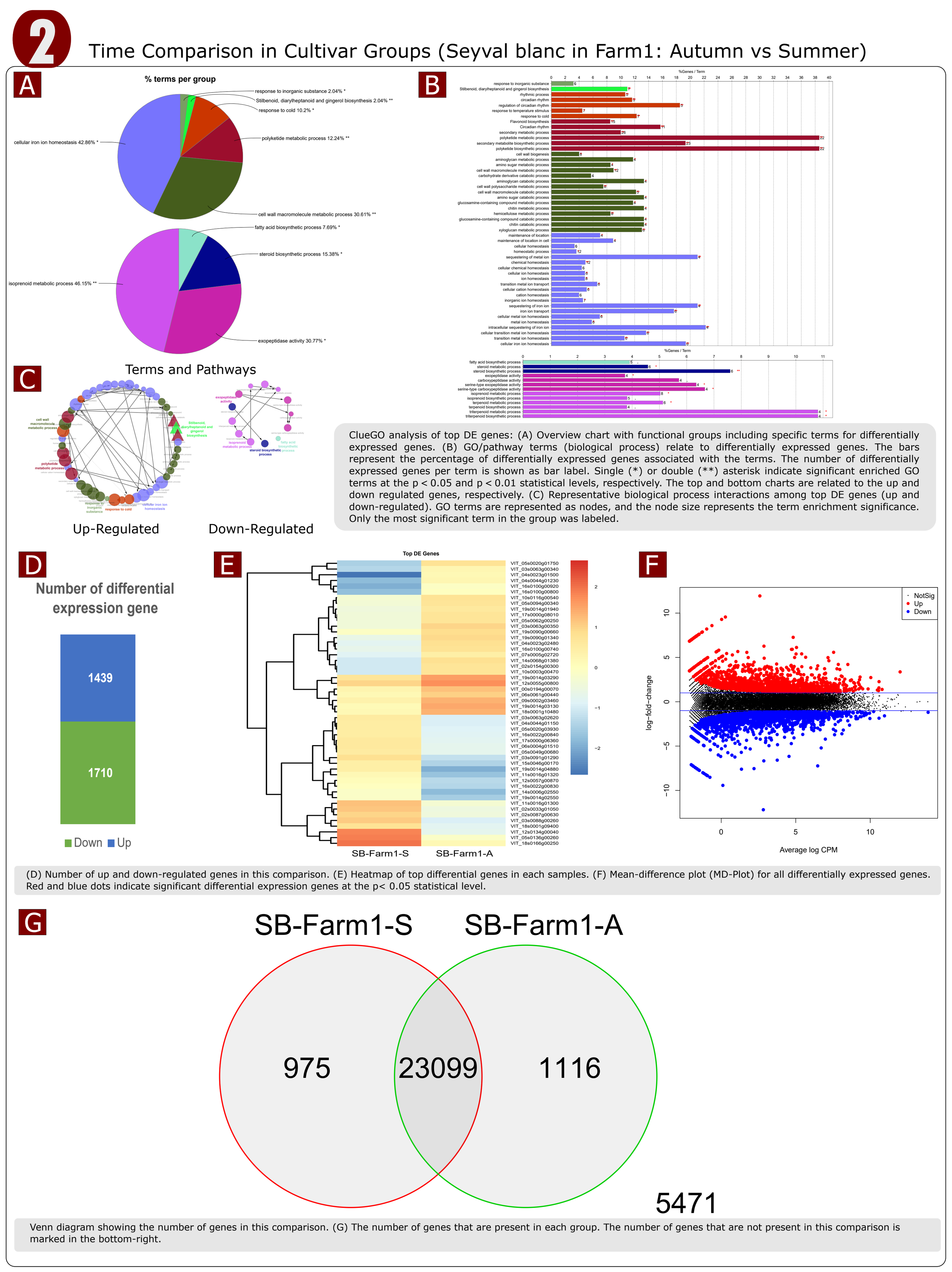


ClueGO analysis of top DE genes: (A) Overview chart with functional groups including specific terms for differentially expressed genes. (B) GO/pathway terms (biological process) relate to differentially expressed genes. The bars represent the percentage of differentially expressed genes associated with the terms. The number of differentially expressed genes per term is shown as bar label. Single (*) or double (**) asterisk indicate significant enriched GO terms at the p < 0.05 and p < 0.01 statistical levels, respectively. The top and bottom charts are related to the up and down regulated genes, respectively. (C) Representative biological process interactions among top DE genes (up and down-regulated). GO terms are represented as nodes, and the node size represents the term enrichment significance. Only the most significant term in the group was labeled.

(D) Number of up and down-regulated genes in this comparison. (E) Heatmap of top differential genes in each samples. (F) Mean-difference plot (MD-Plot) for all differentially expressed genes. Red and blue dots indicate significant differential expression genes at the p< 0.05 statistical level.

Venn diagram showing the number of genes in this comparison. (G) The number of genes that are present in each group. The number of genes that are not present in this comparison is marked in the bottom-right.

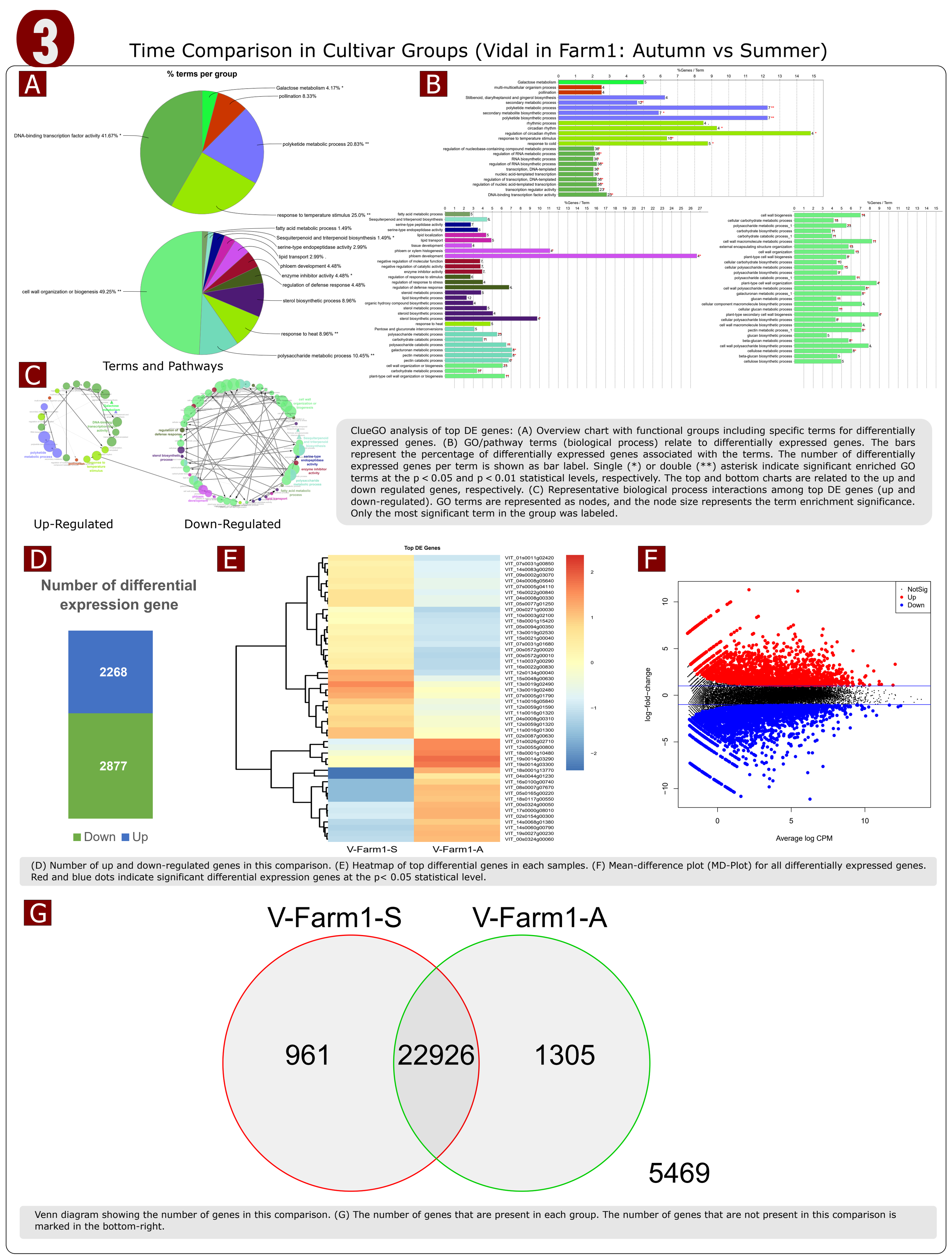


ClueGO analysis of top DE genes: (A) Overview chart with functional groups including specific terms for differentially expressed genes. (B) GO/pathway terms (biological process) relate to differentially expressed genes. The bars represent the percentage of differentially expressed genes associated with the terms. The number of differentially expressed genes per term is shown as bar label. Single (*) or double (**) asterisk indicate significant enriched GO terms at the p < 0.05 and p < 0.01 statistical levels, respectively. The top and bottom charts are related to the up and down regulated genes, respectively. (C) Representative biological process interactions among top DE genes (up and down-regulated). GO terms are represented as nodes, and the node size represents the term enrichment significance. Only the most significant term in the group was labeled.

(D) Number of up and down-regulated genes in this comparison. (E) Heatmap of top differential genes in each samples. (F) Mean-difference plot (MD-Plot) for all differentially expressed genes. Red and blue dots indicate significant differential expression genes at the p< 0.05 statistical level.

Venn diagram showing the number of genes in this comparison. (G) The number of genes that are present in each group. The number of genes that are not present in this comparison is marked in the bottom-right.

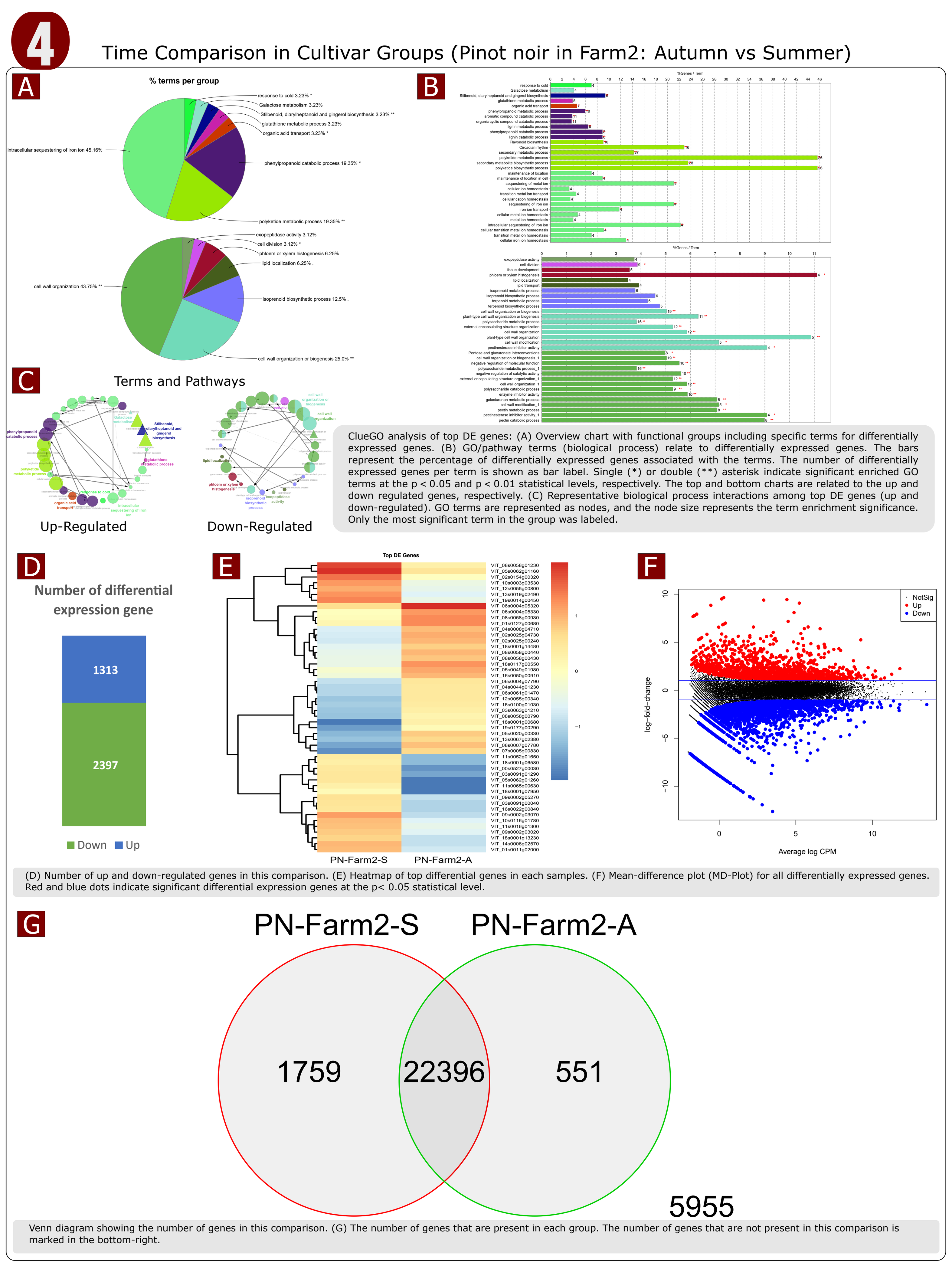


ClueGO analysis of top DE genes: (A) Overview chart with functional groups including specific terms for differentially expressed genes. (B) GO/pathway terms (biological process) relate to differentially expressed genes. The bars represent the percentage of differentially expressed genes associated with the terms. The number of differentially expressed genes per term is shown as bar label. Single (*) or double (**) asterisk indicate significant enriched GO terms at the p < 0.05 and p < 0.01 statistical levels, respectively. The top and bottom charts are related to the up and down regulated genes, respectively. (C) Representative biological process interactions among top DE genes (up and down-regulated). GO terms are represented as nodes, and the node size represents the term enrichment significance. Only the most significant term in the group was labeled.

(D) Number of up and down-regulated genes in this comparison. (E) Heatmap of top differential genes in each samples. (F) Mean-difference plot (MD-Plot) for all differentially expressed genes. Red and blue dots indicate significant differential expression genes at the p< 0.05 statistical level.

Venn diagram showing the number of genes in this comparison. (G) The number of genes that are present in each group. The number of genes that are not present in this comparison is marked in the bottom-right.

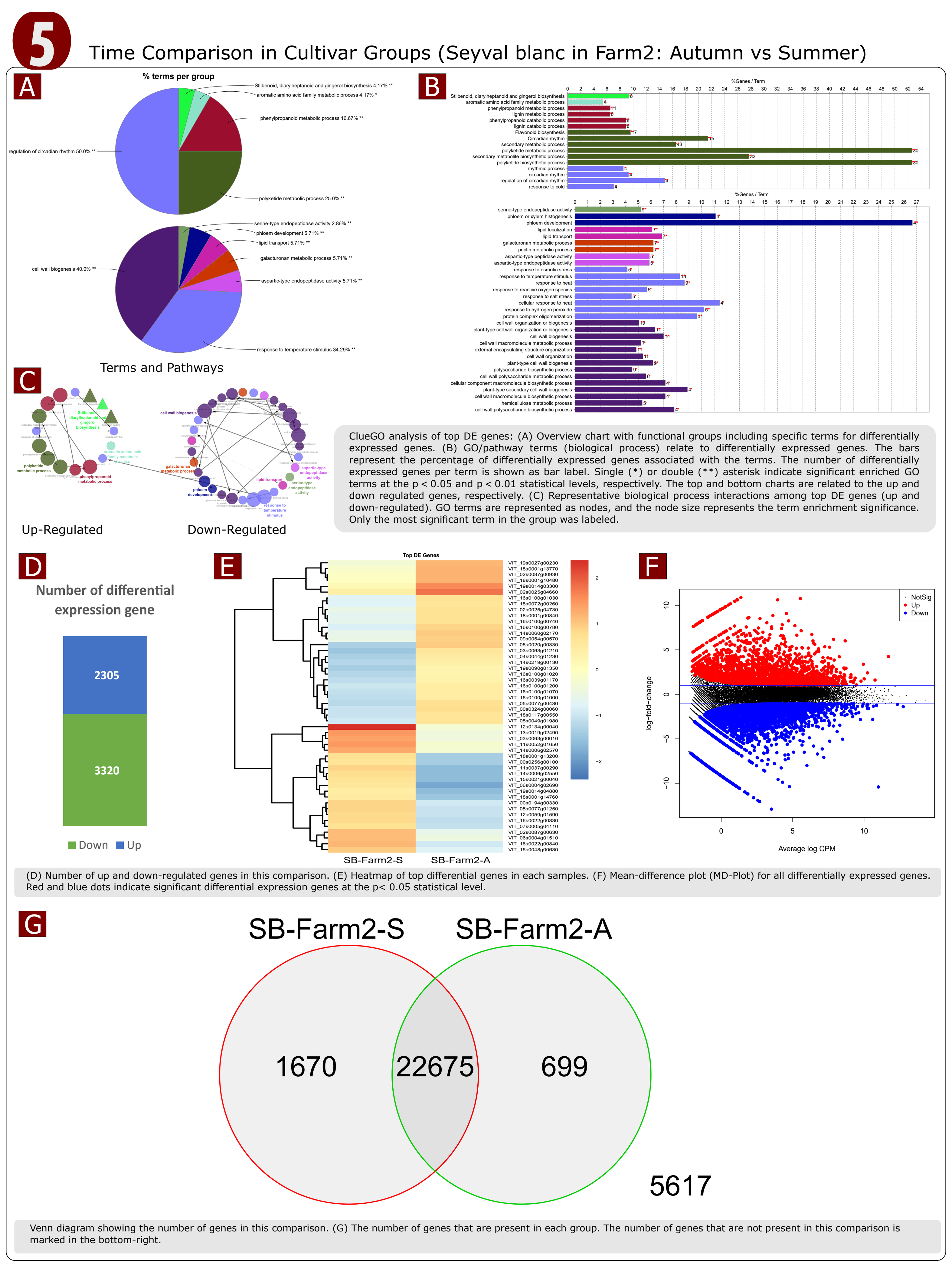


ClueGO analysis of top DE genes: (A) Overview chart with functional groups including specific terms for differentially expressed genes. (B) GO/pathway terms (biological process) relate to differentially expressed genes. The bars represent the percentage of differentially expressed genes associated with the terms. The number of differentially expressed genes per term is shown as bar label. Single (*) or double (**) asterisk indicate significant enriched GO terms at the p < 0.05 and p < 0.01 statistical levels, respectively. The top and bottom charts are related to the up and down regulated genes, respectively. (C) Representative biological process interactions among top DE genes (up and down-regulated). GO terms are represented as nodes, and the node size represents the term enrichment significance. Only the most significant term in the group was labeled.

(D) Number of up and down-regulated genes in this comparison. (E) Heatmap of top differential genes in each samples. (F) Mean-difference plot (MD-Plot) for all differentially expressed genes. Red and blue dots indicate significant differential expression genes at the p< 0.05 statistical level.

Venn diagram showing the number of genes in this comparison. (G) The number of genes that are present in each group. The number of genes that are not present in this comparison is marked in the bottom-right.

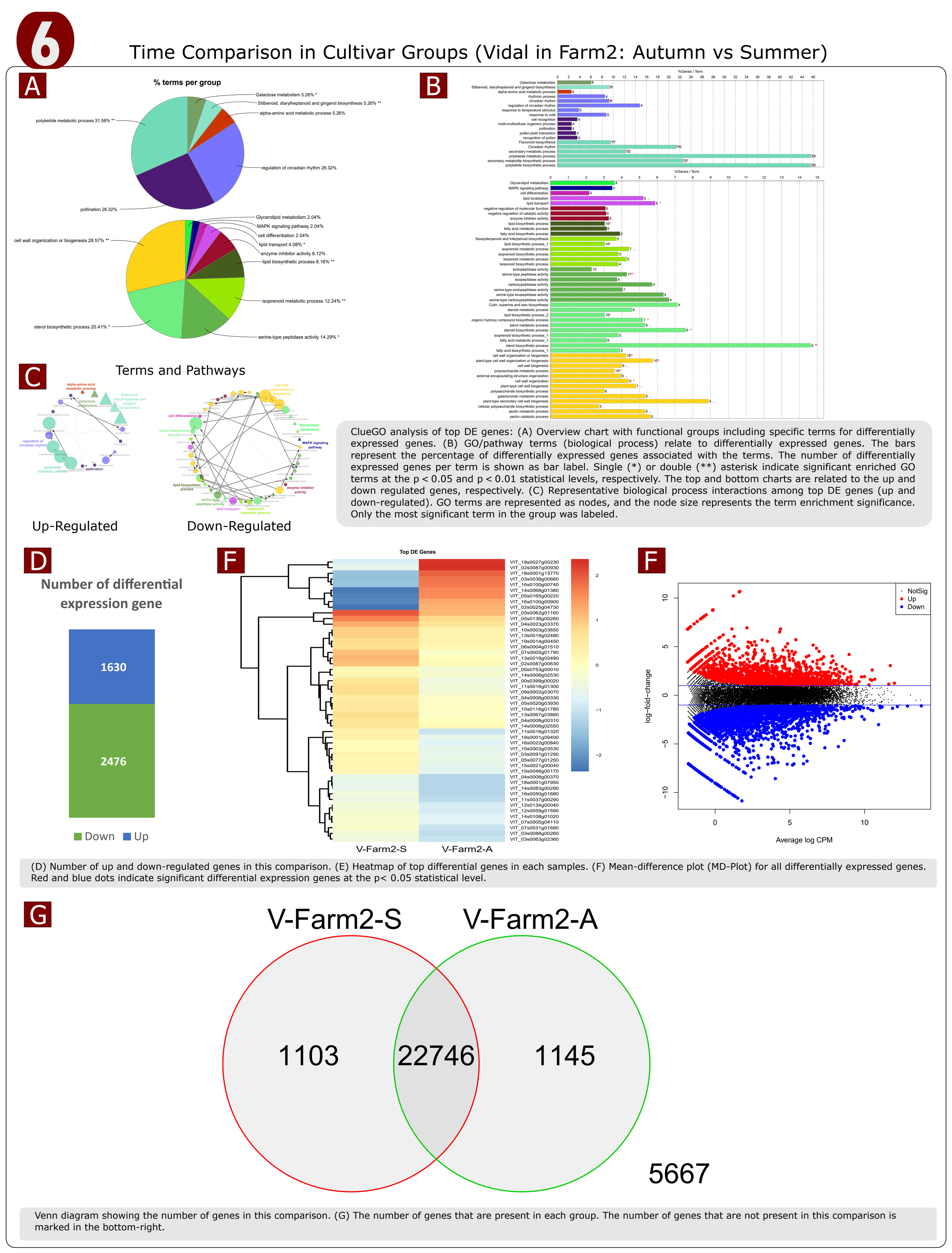


ClueGO analysis of top DE genes: (A) Overview chart with functional groups including specific terms for differentially expressed genes. (B) GO/pathway terms (biological process) relate to differentially expressed genes. The bars represent the percentage of differentially expressed genes associated with the terms. The number of differentially expressed genes per term is shown as bar label. Single (*) or double (**) asterisk indicate significant enriched GO terms at the p < 0.05 and p < 0.01 statistical levels, respectively. The top and bottom charts are related to the up and down regulated genes, respectively. (C) Representative biological process interactions among top DE genes (up and down-regulated). GO terms are represented as nodes, and the node size represents the term enrichment significance. Only the most significant term in the group was labeled.

(D) Number of up and down-regulated genes in this comparison. (E) Heatmap of top differential genes in each samples. (F) Mean-difference plot (MD-Plot) for all differentially expressed genes. Red and blue dots indicate significant differential expression genes at the p< 0.05 statistical level.

Venn diagram showing the number of genes in this comparison. (G) The number of genes that are present in each group. The number of genes that are not present in this comparison is marked in the bottom-right.
